# Microtubules coordinate with actin assembly and disassembly to elongate cortical actin cables

**DOI:** 10.64898/2026.11.03.762817

**Authors:** Wei-Chien Chou, Brooke M. McCartney

## Abstract

The coordination of actin filament elongation, bundling and turnover within bundled actin structures is poorly understood. Nurse cells extend bundles of actin filaments (cables) from the cortex during Drosophila oogenesis. We previously showed that cable assembly requires a developmentally-regulated microtubule (MT) network. Here we demonstrate that MTs coordinate with actin filament assembly and disassembly to promote the persistent elongation of actin cables. Disrupting the MT network caused actin cables to detach from the cortex and become free in the cytoplasm. The subsequent fate of those free cables, shrinking over time or maintaining a stable length, depended on the MT manipulation. Furthermore, maintaining cortical continuity and persistent cable elongation requires a regulated imbalance between filament assembly and disassembly. Favoring disassembly caused cable detachment and shrinkage while reducing disassembly allowed free cables to continue elongating in the cytoplasm. Our results also suggest that both filament assembly and disassembly occur at the cortical filament barbed-ends protected from excessive disassembly by Capping Protein.

**Summary:** This study demonstrates that actin cable elongation from the cortex to the nucleus during Drosophila oogenesis requires imbalanced actin filament assembly and disassembly at the cortical filament barbed-ends and regulation by the coaligned microtubule network.

## Introduction

Actin filaments are organized into a remarkable variety of networks with varying geometries that are critical for many basic cellular functions, including cell migration, mechanosensing, adhesion, endocytosis, and cytokinesis. Branched filament networks and linear filament bundles underlie this spatial diversity (Blanchoin et al., 2014). Actin filament bundles are collections of crosslinked, linear filaments that form the core of a variety of structures, including filopodia, microvilli, stereocilia, stress fibers, and the cell cortex. Parallel bundles are composed of filaments aligned with uniform polarity: all barbed, fast-growing ends of the filaments are at one end of the bundle while the pointed, slow growing ends are at the other (Rajan et al. 2023). The number and length of actin filaments within parallel actin bundles are highly regulated and correlate with specific functional demands. For example, filopodia at the leading edge of motile cells contain approximately 10 filaments and grow to roughly 10 µm (Mogilner and Rubinstein, 2005), while a 10 µm stereocilium contains thousands of filaments (Tilney et al., 1980). Drosophila bristles, mechanosensing organs containing multiple bundles of hundreds of filaments, can reach 300 µm in length (Tilney et al., 1995). The organization, structure, and mechanical properties of actin bundles are dictated in part by actin filament bundling proteins. More than 10 families of actin bundling proteins have been identified, and bundles are typically crosslinked by two or more actin bundlers that can have both partially redundant functions and unique properties (Rajan et al. 2023; Michelot and Drubin, 2011; Blanchoin et al., 2014; Lappalainen, 2016; Pollard, 2016; Sherer and Courtemanche, 2022). While actin bundles are produced from a common set of highly conserved actin assembly factors and bundling proteins, the mechanisms that coordinate filament assembly, bundle formation and bundle maintenance are poorly understood.

Short-lived bundles like filopodia can have a turnover time on the order of minutes and are disassembled through the “demolition pathway” that includes proteins like Cofilin, Aip1, Twinfilin and Capping protein (Goode at al. 2023). Long-lived actin bundles like those in microvilli and stereocilia have turnover times ranging from hours to days and utilize the “dynamic maintenance pathway” to balance disassembly with assembly and promote the persistence of the bundle (Goode et al., 2023). Two modes of filament dynamic maintenance in more stable bundles have been identified: (1) treadmilling, with balanced barbed-end assembly and pointed-end disassembly, as in microvilli (Tyska and Mooseker, 2002; Meenderink et al., 2019; Kachar, 2019) and (2) barbed-end dynamic maintenance where balanced assembly and disassembly both occur at the filament barbed-ends, as in stereocilia (Drummond et al., 2015; Narayana et al., 2015). The role of filament bundling in general and individual bundlers specifically in regulating demolition or dynamic maintenance is unclear. In some contexts, bundling proteins appear to enhance filament disassembly (Brietsprecher et al., 2011; Kemp Jr. and Brieher, 2018), while in others, bundling suppresses disassembly (Tilney et al., 2003; Chikireddy et al., 2024).

The elongating actin cables found in the nurse cells (NCs) of the Drosophila ovary are a valuable in vivo model for dissecting mechanisms of actin bundle assembly and regulation. A single *Drosophila* egg chamber, or follicle, within the ovary contains a syncytium of 16 germ cells connected by ring canals and surrounded by a somatic epithelium of follicle cells (Fig. 1A). A posterior germ cell becomes the oocyte while the other 15 become nurse cells that deliver mRNAs, proteins, and organelles to the growing oocyte via the ring canals. At a late stage of oogenesis (stage 11; s11), the nurse cells expel their remaining cytoplasm into the oocyte, roughly doubling its volume in approximately 100 min (Fig. 1A; Alsous et al. 2021). This cytoplasmic dumping is driven by an initial phase of hydraulic transport supported by baseline cortical tension and a second phase characterized by actomyosin contractions of the NC cortex (Alsous et al. 2021). Massive actin cable arrays initiate and elongate during s10B (Fig. 1A) and we defined 4 substages (s10B.1, s10B.2, s10B.3 and s10B.4) to facilitate the analysis of actin cable assembly (Chou et al., 2026). These cables appear to enable s11 cytoplasmic dumping by pushing and rolling the nurse cell nuclei away from the ring canals (Huelsmann et al., 2013; Logan et al., 2022). Without the actin cables, nuclear obstruction of the ring canals blocks the completion of oogenesis (Cant et al., 1994; Mahajan-Miklos and Cooley, 1994; Logan et al., 2022; Gates et al., 2009).

**Figure 1:**
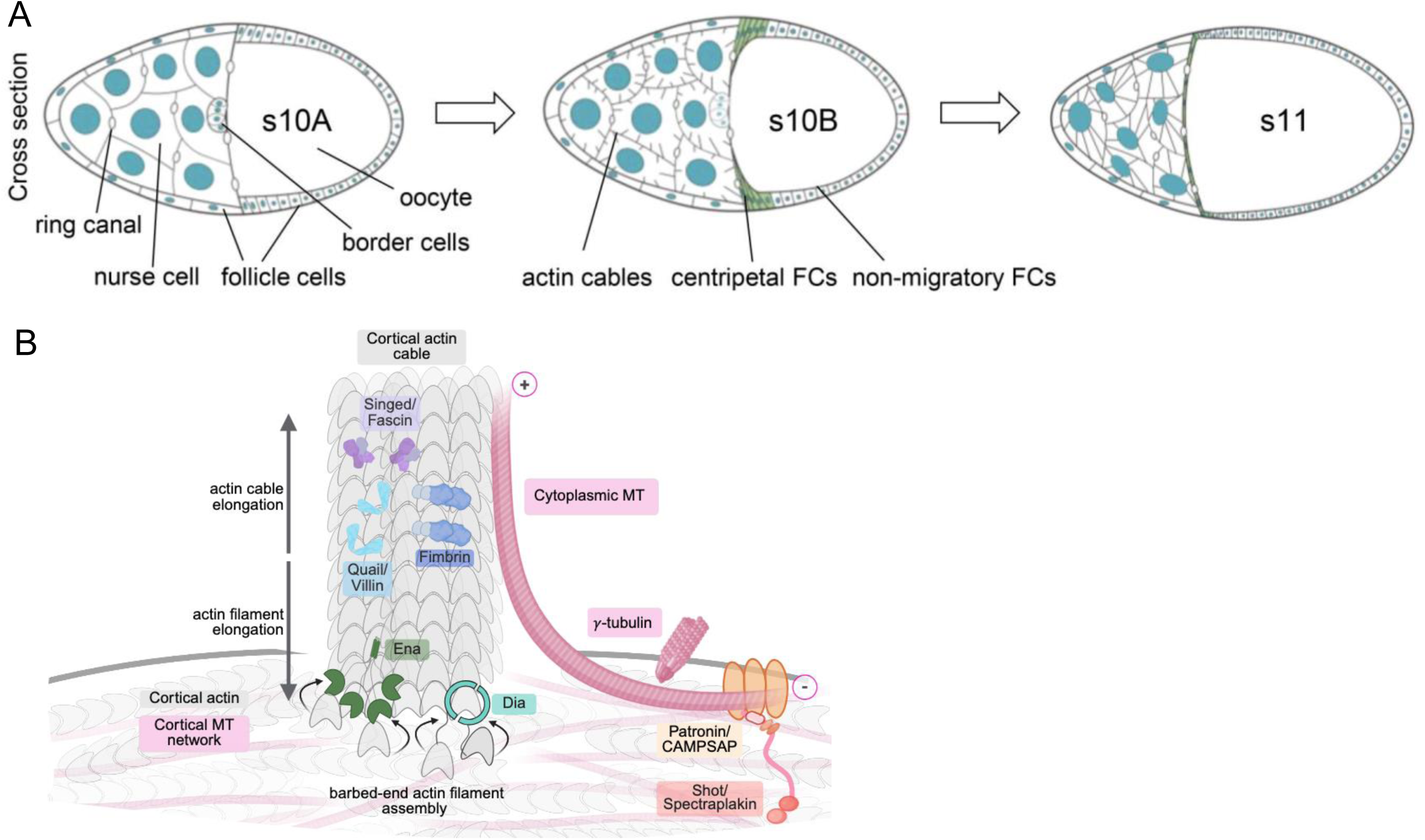
Schematic representations of oogenesis during s10-11 and our current understanding of mechanisms regulating actin cable assembly and the Ace-MT network. (A) Schematic of egg chambers during s10-11 of oogenesis (from Fig. 1A, Chou et al., 2026). Ring canals connect the nurse cells to each other and to the oocyte and enable cytoplasmic transport into the growing oocyte. During s10B, actin cables initiate at the actin cortex and elongate toward the nurse cell nuclei. The actin cables reposition the nurse cell nuclei away from the ring canals to enable rapid cytoplasmic transport (cytoplasmic “dumping”) in s11. (B) During s10B, Dia and Ena elongate the barbed-ends of actin filaments at the cortex thereby pushing the entire cable away from the cortex and toward the nucleus. Actin filaments are bundled into cables by Singed/Fascin, Quail/Villin and Fimbrin. The Ace-MT network is nucleated at the cortex by γ-tubulin and Ace-MTs are tethered to the cortex via Patronin/CAMSAP and Shot/Spectraplakin. Ace-MTs lie parallel to the cortex and extend into the cytoplasm where they coalign with actin cables and co-regulate (Chou et al., 2026). (modified from Fig. 9B Chou et al., 2026) Created in Biorender.

Nurse cell actin cables can elongate up to 30 µm and are composed of 2-4 μm unit bundles, each of which contains approximately 26 parallel actin filaments (Guild et al., 1997). In mature cables, the unit bundles appear to be bundled in a staggered, overlapping pattern. All unit bundles have their barbed-ends oriented toward the cortex with the barbed-ends of the most proximal bundle at the cortex (Fig. 1B; Guild et al., 1997). We and others have previously shown that actin filament assembly is mediated by cortically localized Diaphanous (Dia) and Enabled (Ena), and that bundling requires Quail/Villin, Singed/Fascin, and Fimbrin (Fig. 1B; Cant et al., 1994; Matova et al., 1999; Gates, 2009; Huelsmann et al., 2013; Logan et al., 2022; Chou et al., 2026). We recently demonstrated that a developmentally-regulated, stable and acetylated microtubule (Ace-MT) network is required for actin cable assembly (Chou et al., 2026). These Ace-MTs are nucleated by γ-tubulin, cortically-tethered via Shot and Patronin, and strongly coalign with the elongating actin cables (Fig. 1B). Depleting the Ace-MT network resulted in fewer, slower growing cables (Chou et al., 2026). We also demonstrated that the density of the total MT network and the Ace-MT network is comparable throughout s10B, suggesting that all, or virtually all MTs, are acetylated at this stage (Chou et al., 2026). We will refer to the Ace-MTs as simply MTs in this work.

Here, we show that MTs play an unexpected and essential role in maintaining the cortical continuity of the actin cables. Disrupting the MT network frequently caused the remaining elongating cables to detach from the cortex and become free in the cytoplasm. These free cables either maintained a stable length or shrank, depending on the MT manipulation. Surprisingly, our results further suggest that the regulated disassembly of actin filaments is required for the cortical continuity and continued assembly of actin cables. Directly biasing the process toward filament assembly or toward filament disassembly produced free cables that predictably elongated or shrank. Our results indicate that both filament assembly and disassembly occur at the filament barbed ends. Together, our data suggest that MTs promote the barbed-end dynamics of actin filaments that are required to maintain the cortical association and persistent elongation of the nurse cell actin cables that are essential for oogenesis.

## Results

### CLIP-190 and EB1 are required for the MT network and to promote cortical actin cable assembly

To begin to understand how MTs might impact actin cable assembly, we examined the roles of CLIP-190 (Drosophila CLIP-170, CLIP) and EB1. These proteins can interact with the actin assembly factor Dia, facilitate actin-MT crosslinking, and/or modulate MT dynamics in vitro and in cultured cells (Folker et al., 2005; Lewkowicz et al., 2008; Komarova et al., 2009; Henty-Ridilla et al., 2016). We reduced CLIP or EB1 in nurse cells using *UAS-dsRNA* expression driven by a strong, germline GAL4 driver (*MTD>*). For all genetic manipulations presented here, representative phenotypes were reproduced using two independent approaches per gene and KD using at least one approach per gene was demonstrated by RT-PCR or immunolabeling unless previously described (Table 1 and Fig. S1). In CLIP and EB1 KDs, we observed a significant loss of both the cortical and cytoplasmic MT networks (Fig. 2A-C,E,H), consistent with their roles in regulating MT dynamics and growth in vitro and in cells (Perez et al., 1999; Piehl and Cassimeris, 2002; Bieling et al., 2008; Maurer et al., 2014). Given the dramatic loss of MTs, it was not surprising that CLIP or EB1 KD caused a significant reduction in actin cable density and length (Fig. 2A-C, F, G) throughout s10B. Shorter cables are consistent with a slower elongation rate (Chou et al., 2026). In addition to the overall loss of actin cables, we were surprised to find that roughly 65-75% of the cables that remained were free in the cytoplasm and often clustered around the nucleus (Fig. 2B,C yellow arrowheads, I). In wild-type (WT) nurse cells, actin cables are primarily associated with the cortex where Dia and Ena assemble actin filaments (Fig. 2A; Gates et al., 2009; Huelsmann et al., 2013; Logan et al., 2022), but we found a small population of free cables in the cytoplasm of control (WT or driver control *MTD>*) nurse cells (Fig. 2I). Such free cables have not been previously reported. If free cables resulted from depleting the MT network, we predicted that we should also see free cables in nurse cells that have lost their MTs due to γ-tubulin, Patonin, or Shot KD. We previously showed that reduction in any of these resulted in a significant loss of MTs and actin cables (Chou et al, 2026 and Fig. 2D, E-H). Consistent with that hypothesis, we observed a significant increase in the fraction of free cables in γ-tubulin, Patonin, and Shot KDs (approx. 50-55%; Fig. 2D,I, Fig. S2A-C) and these also tended to cluster around the nucleus (Fig. 2D and Fig. S2A,B, yellow arrowheads).

**Figure 2:**
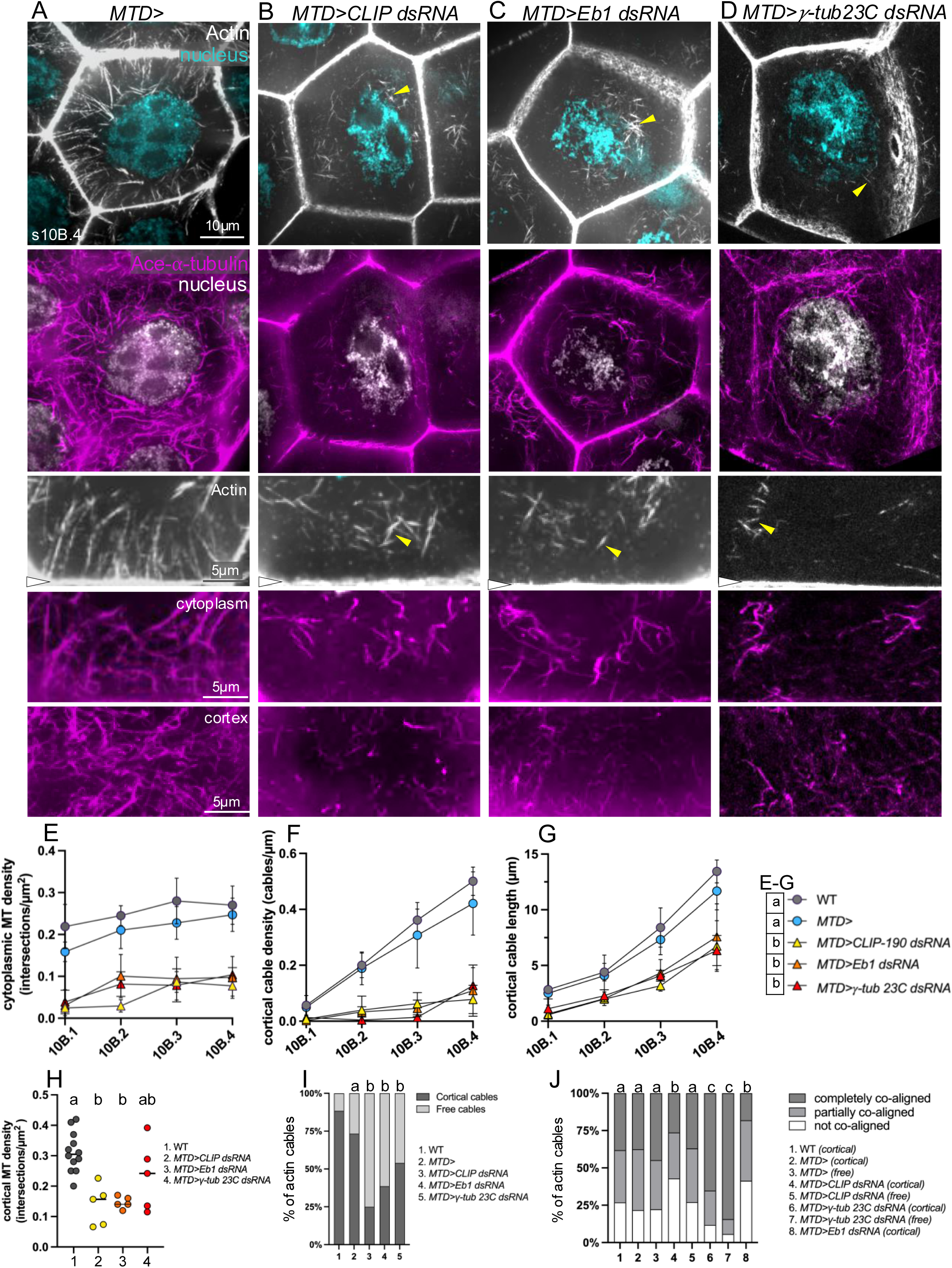
CLIP, EB1 and γ-tubulin regulate the Ace-MT network and control cortical actin cable assembly. (A-D) Representative images of fixed s10B.4 nurse cells triple-labeled for actin cables (grey; phallodin), DNA (DAPI; grey or cyan) and the Ace-MT network (magenta; anti-Ace-α-tubulin) from (A) control *MTD>* (B) *MTD>CLIP dsRNA,* (C) *MTD>Eb1 dsRNA*, and (D) *MTD>γ-tubulin 23C dsRNA* genotypes. These are shown at lower magnification (scale bar = 10 µm) and higher magnification (scale bar = 5 µm). Images were taken of the same nurse cells for the lower magnification views and the higher magnification views of actin and cytoplasmic Ace-MTs. Independent, stage-matched nurse cells were used for the images of the cortical Ace-MT network. The cortical Ace-MT network is shown *en face* and all other images are cross-sections. Yellow arrowheads indicate free cables, white arrowheads mark the cortex. (E) Quantification of cytoplasmic Ace-MT density across s10B in wild type nurse cells (WT), driver control nurse cells (*MTD>*) and nurse cells upon depletion of CLIP, EB1, or γ-tubulin. E-G share a key. Two-way ANOVA with Tukey’s multiple comparison test. Conditions sharing letters are not significantly different from one another, conditions with different letters are significantly different, p<0.0001, n=7-12 per substage. (F,G) Quantification of cortical actin cable density (F) and cortical actin cable length (G) across s10B in wild type nurse cells (WT), driver control nurse cells (*MTD>*) and nurse cells upon depletion of CLIP, EB1, or γ-tubulin. E-G share a key. Two-way ANOVA with Tukey’s multiple comparison test. Conditions sharing letters are not significantly different from one another, conditions with different letters are significantly different, p<0.0001, n=7-12 per substage. (H) Quantification of cortical Ace-MT density at s10B.3 in genotypes as shown. The *MTD>Eb1 dsRNA* symbols and line are obscured by that of *MTD>CLIP dsRNA*. Two-way ANOVA with Tukey’s multiple comparison test. Conditions sharing letters are not significantly different from one another, conditions with different letters are significantly different, p<0.005, n=5-12 per genotype. (I) Quantification of the proportion of cortical versus free cables during s10B in genotypes as shown. All MT-depleted genotypes show a significant increase in the fraction of free cables. Chi-square test for goodness fit, (2) is expected compared to (3-5) observed. Conditions sharing letters are not significantly different from one another, conditions with different letters are significantly different p<0.0001, n=20-45 egg chambers per genotype with all substages represented. (J) Quantification of co-alignment between cortical or free actin cables and Ace-MTs in s10B.4 in genotypes as shown. Chi-square test for goodness fit, conditions sharing letters are not significantly different from one another, conditions with different letters are significantly different p<0.0001, n=7-10 egg chambers per genotype, 200-450 cables/genotype were analyzed.

**Table 1:** Summary of protein depletion/knockdown efficiency and validation of phenotypes with additional genotypes.

### Free cables initiate at the cortex and detach over time

Free cables could result from ectopic cable assembly in the cytoplasm, or from the detachment of cables from the cortex. To distinguish between those alternatives, we labeled live egg chambers with Sir-actin or SPY-actin (Logan et al., 2022) and tracked actin cable elongation and cortical association for 75-90 min. In control nurse cells (either WT or the *MTD>* driver control), actin cables emerged from the cortex, maintained continuity with the actin cortex, and elongated at approx. 0.12 µm/min as we previously described (Fig. 3A,E arrowheads, H,I; Logan et al., 2022). Although we did see a small number of free cables in fixed, wild-type nurse cells, we did not observe them in live cells. This may be because they are hidden by the large population of elongating cortical cables. In CLIP, EB1 or γ-tubulin KDs, actin cables were initially associated with the cortex but lost their cortical continuity over time (Fig. 3B-D, arrowheads, Fig. S3A,B). While we can’t rule out ectopic cytoplasmic cable assembly, we never observed it.

**Figure 3:**
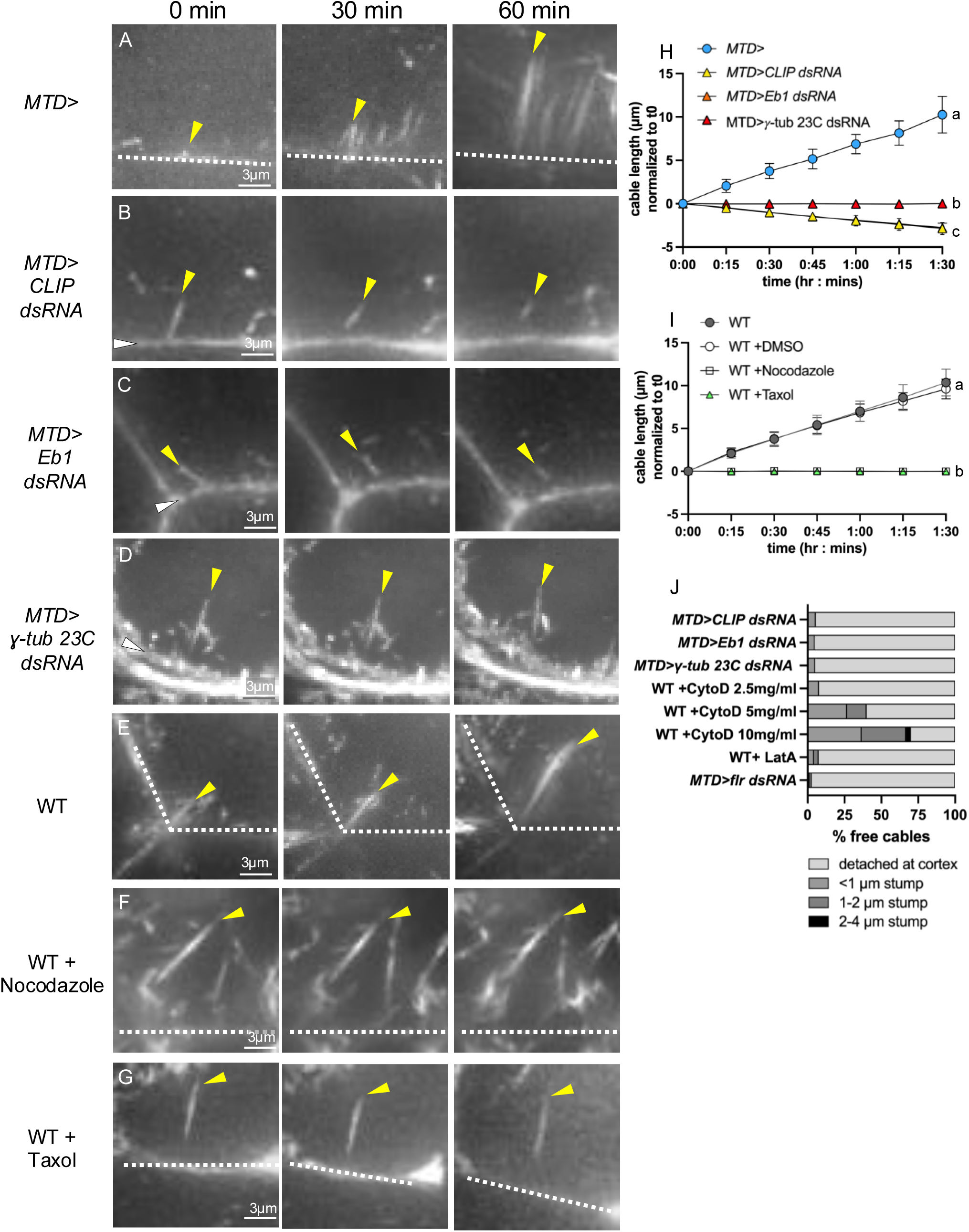
Free cables detach from the cortex and either shrink or maintain a stable length depending on the type of MT network manipulation. (A-G) Images of s10B actin cables in live nurse cells labeled with SPY-Actin in genotypes and conditions as shown (scale bar = 3 µm). Actin cables were imaged for 90 minutes; representative time points at t=0, t=30, and t=60 minutes are shown. The cortex is indicated with white dashed lines or white arrowheads, and representative cables are indicated by yellow arrowheads. (H,I) Quantification of cable length over time in genotypes and conditions as shown. Initial cable lengths were normalized to 0 µm at time = 0:00 for comparison. In WT and *MTD>* driver control cells, elongating cortical cables were measured. In KDs and drug-treated conditions, cables detach at time zero and free cable lengths were measured. Wild-type (E,I) and control (*MTD>*, A,H) cortical cables appear to elongate at a steady rate and remain associated with the cortex. Cortical cables detach upon reduction of CLIP (B) or EB1 (C) and detached cables shrink over time (H). Cortical cables detached and maintained stable length upon reduction of γ-tubulin (D,H) or in cells treated with nocodazole (F,I) or taxol (G,I). Simple linear regression with ANOVA. Genotypes sharing letters are not statistically different from one another. p<0.0001. n=20∼25 cables per genotype/condition. (J) Quantification of detaching vs breaking cables in genotypes and conditions as shown. Only treatment with Cytochalasin D produced significant cable breakage. n=39-94 cables per genotype/condition.

Cables that lose cortical association could be detaching from the cortex, breaking internally, or both. We examined actin cables that were in the process of losing their cortical association in CLIP, EB1 and γ-tubulin KDs and found that >95% of those cables appeared to detach from the cortex and did not leave a resolvable stump (Fig. 3J). In fewer than 5% of cases, we observed stumps <1 µm in length. As a control, we examined cables in WT cells treated with Cytochalasin D (CytoD). At high concentrations, CytoD has been reported to bind to filament sides and induce breakage (Mitani et al., 2025). We found that the fraction of cables that break internally increased significantly with increasing concentrations of CytoD (Fig. 3J). Together, these data suggest that in CLIP, EB1 and γ-tubulin KDs free cables are the result of cables detaching at, or very near, the actin cortex.

We previously showed that the growing cortically associated actin cables coalign with MTs (Chou et al., 2026). Thus, we considered the possibility that the actin cables are losing their cortical continuity due to the loss of cortically tethered, coaligning MTs in the MT depleted nurse cells. In control nurse cells, approx. 75-80% of cortical cables and free cables coaligned at least partially with cytoplasmic MTs in s10B (Fig. 2J and Chou et al., 2026). In CLIP and EB1 KDs, the degree of coalignment between cortical cables and MTs was significantly reduced to approx. 60% (Fig. 2J), but free cable coalignment in CLIP KDs was not significantly different from free cables in control nurse cells (Fig. 2J). Interestingly, in γ-tubulin KDs, coalignment significantly increased such that approx. 90% of both cortical and free cables were partially or completely coaligned with MTs (Fig. 2J; Chou et al., 2026). This suggests that the presence of free cables cannot be simply attributed to the lack of coaligned MTs.

### Different MT network perturbations affect the fate of free cables in different ways

Surprisingly, actin cables that detached from the cortex in CLIP-190 and EB1 KDs shrank throughout the imaging period at a rate of approx. 0.03 µm/min (Fig. 3B,C,H). This suggests that once these cables lose their association with the cortex, the known site of new Dia- and Ena-mediated filament assembly, they disassemble. Depleting the MT network through reduction of γ-tubulin or treatment of wild type nurse cells with 7.5 µg/ml nocodazole at the start of imaging produced a strikingly different outcome; free cables did not shrink but instead maintained a constant length over the same time period (Fig. 3D,F,H,I; the slope of the lines did not significantly deviate from zero). We previously showed that nocodazole treatment significantly reduced the density of the MT network (Chou et al, 2026). This suggests that while the MT network is generally required to maintain the cortical continuity of actin cables, the properties of the remaining MTs may impact the fate of the free cables. For example, in CLIP or EB1 KD nurse cells, the remaining MTs are depleted of at least one associated protein. In the γ-tubulin KD or nocodazole treated nurse cells, we predict that the remaining MTs would still be associated with their typical binding proteins.

We previously reported that actin cables detach from the cortex in nurse cells labeled with high concentrations of Tubulin Tracker, a taxol derivative, or treated with MT-stabilizing taxol itself (Logan and McCartney, 2020). We confirmed that actin cables detach from the cortex in WT, taxol-treated nurse cells (Fig. 3G, Fig. S3C), and found that these detached cables maintained a constant length over time, as we observed with nocodazole treatment or γ-tubulin KD (Fig. 3G,I). This suggests that stabilizing the MT network, at least with taxol, can also cause loss of cortical cables that appear to remain stable in the cytoplasm.

Taken together, these data support the model that the MT network is required for the cortical continuity of actin cables. Furthermore, our data suggest that specific properties of the MTs (e.g. structural, mechanical, dynamic, binding protein association) may determine whether the free cables shrink or maintain a constant length in the cytoplasm.

### Decreased cable elongation rate or altered actin filament bundling does not affect actin cable cortical continuity

How do actin cables maintain cortical continuity while elongating? Our current model of actin cable assembly is that cortical Dia and Ena collaborate to elongate the barbed ends of actin filaments at the cortex that are bundled into unit bundles and cables by Quail/Villin, Singed/Fascin, and Fimbrin (Fig. 1B; Guild et a., 1997; Logan et al., 2022; Chou et al. 2026). We predicted that cortical continuity may be maintained by the persistent actin filament elongation mediated by Ena and Dia. We previously showed that reduction of either Ena or Dia significantly reduced actin cable density and elongation rate (Logan et al., 2022; Chou et al., 2026) and decreased MT tethering to the cortex but did not impact the degree of actin cable-MT coalignment (Chou et al, 2026). To test the role of the normal filament elongation rate in maintaining cortical continuity, we examined Dia and Ena KDs for the presence of free cables. Of the remaining cables in Dia and Ena KDs, the fraction of free cables was indistinguishable from the control (Fig. 4A-C,G).

**Figure 4:**
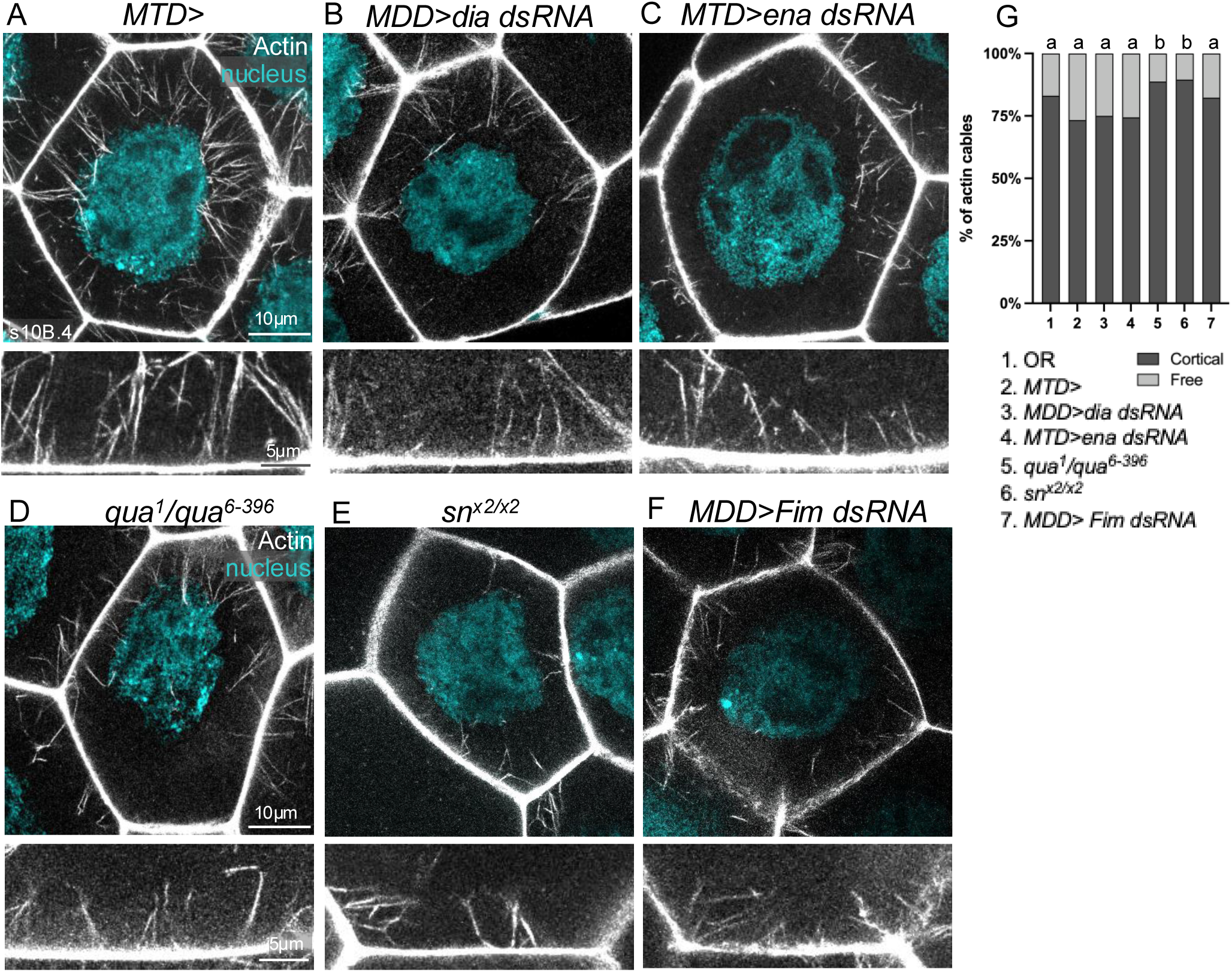
Neither reduction in the rate of actin filament assembly nor disruption of actin filament bundling affects the cortical continuity of actin cables. (A-F) Representative images of fixed s10B.4 nurse cells in (A) control *MTD>* (B) *MDD>dia dsRNA* (C) *MTD>ena dsRNA* (D) *qua^1^/qua^6-396^*(E) *sn^x2/x2^* and (F) *MDD>Fim dsRNA* nurse cells. The nurse cells were double labeled to reveal actin cables (phalloidin, grey) and nuclei (DAPI, cyan) and are shown in cross section at lower magnification (scale bar = 10 µm) and higher magnification (scale bar = 5 µm). (G) Quantification of the proportion of cortical versus free cables during s10B. Reduction of Dia, Ena, or Fimbrin did not result in a significant increase of free cables, but reduction of Quail or Singed surprisingly increased the cortical association of remaining cables. Chi-square test for goodness fit, Conditions sharing letters are not significantly different from one another, conditions with different letters are significantly different p<0.0001, n=20-45 egg chambers per genotype.

We previously showed that nurse cells depleted of the filament bundlers Quail/Villin, Singed/Fascin, or Fimbrin all had a similar reduction in cable number and cable length compared to Dia and Ena, but that the MT network was significantly altered with reduced cytoplasmic and cortical MT density, shorter MTs, and significantly enhanced coalignment between the remaining actin cables and MTs (Chou et al., 2026). The cortical continuity of the remaining cables in all filament bundling mutants/KDs was maintained and even somewhat enhanced compared to control (Fig. 4D-G). Thus, loss of individual actin filament bundlers did not reduce the cortical continuity of the remaining actin cables.

### Modulating actin filament assembly and disassembly reduced cortical continuity and determined the fate of free cables

The dynamics of actin filament assembly and disassembly have been well characterized in vitro and in simplified cellular systems, but dynamics within in vivo filament bundles is poorly understood. The persistent actin bundles in microvilli and stereocilia require continuous filament remodeling in the form of balanced assembly and disassembly to maintain their stable length (Tyska and Mooseker, 2002; Meenderink et al., 2019; Narayanan et al., 2015; Drummond et al., 2015). Because nurse cell cortical actin cables are actively elongating, any potential disassembly must necessarily be exceeded by assembly and thus imbalanced. To determine whether filament disassembly occurs in these actin cables and whether it impacts cortical continuity and elongation, we altered the degree of imbalance between assembly and disassembly. These changes generated significant numbers of free cables and had predictable effects on free cable fate.

We examined cables under conditions that favor disassembly in two ways: indirectly with Latrunculin A (LatA) and directly through reduction of LIMK. LatA binds to free monomers and reduces the pool available for filament assembly (Coué et al., 1987) thereby blocking assembly and indirectly favoring any disassembly that may be occurring. Although decreasing the cable elongation rate by reduced Ena or Dia did not result in cable detachment (Fig. 4G), LatA treatment caused a dramatic increase in free cables (Fig. 5A,F, Fig. 6B, Fig. S3D), and we showed that approx. 92% of those cables detached from the cortex rather than breaking internally (Fig. 3J). This suggests that continuous assembly is required to maintain the cortical continuity of the actin cables. Furthermore, we found that LatA-induced free cables shrank at a rate of approx. 0.04 µm/min (Fig. 5G), similar to the free cables in CLIP or EB1 KDs (Fig. 3H). This is consistent with the model that under some conditions free cables require new subunit addition to maintain themselves in the cytoplasm and without it, the free cables shrink due to disassembly.

**Figure 5:**
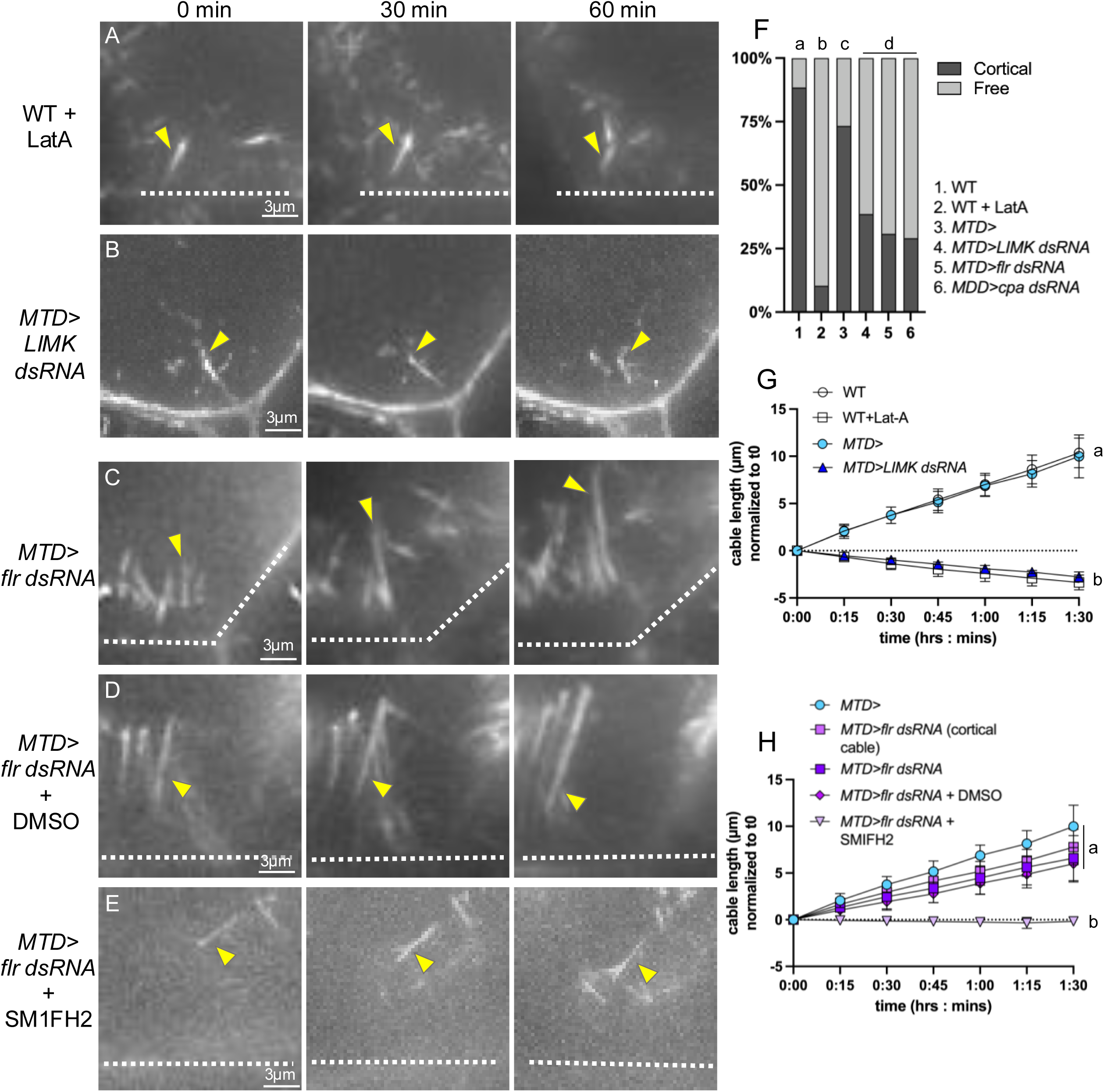
Promoting either actin filament disassembly or actin filament assembly produced free cables that shrank or grew respectively. (A-E) Images of s10B actin cables in live nurse cells labeled with SPY-actin. Actin cables were imaged for 90 minutes; representative time points at t=0, t=30, and t=60 minutes are shown. The cortex is indicated with white dashed lines or white arrowheads, and a representative cable is indicated with yellow arrowheads. (scale bar = 3 µm) (F) Quantification of the proportion of cortical versus free cables across s10B. Chi-square test for goodness fit, Conditions sharing letters are not significantly different from one another, conditions with different letters are significantly different, p<0.0001, n=25-50 egg chambers per genotype. (G,H) Quantification of cable length over time for genotypes and conditions shown. Initial cable lengths were normalized to 0 µm at time = 0:00 for comparison. In wild type and *MTD>* control cells, elongating cortical cables were measured. In knockdowns and drug-treated conditions, cables detach at time zero and free cable lengths were measured. Genotypes sharing letters are not statistically different from one another. p<0.0001. Simple linear regression with ANOVA. n=20∼25 cables per genotype/condition.

**Figure 6:**
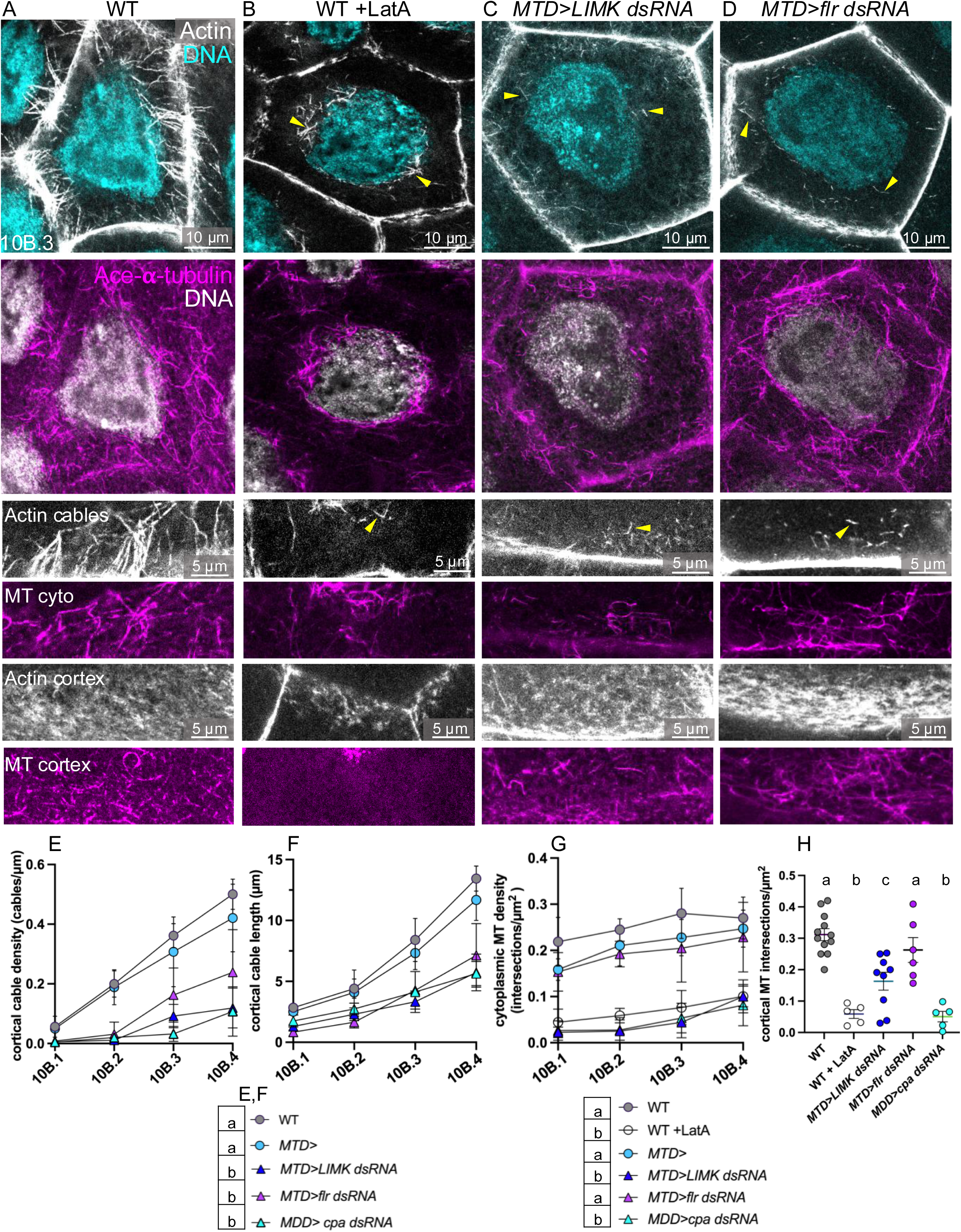
Manipulation of filament assembly and disassembly results in free cables and other changes in actin and MT networks. (A-D) Representative images of fixed s10B.3 nurse cells triple-labeled for actin cables (grey; phallodin), DNA (DAPI; grey or cyan) and the Ace-MT network (magenta; anti-Ace-α-tubulin) in (A) WT, (B) WT + LatA, (C) *MTD>LIMK dsRNA,* and (D) *MTD>flr dsRNA* nurse cells. These are shown at lower magnification (scale bar = 10 µm) and higher magnification (scale bar = 5 µm). Images were taken of the same nurse cells for the lower magnification views and the higher magnification views of actin and cytoplasmic Ace-MTs. Independent, stage-matched nurse cells were used for the images of the actin cortex and cortical Ace-MT network. The cortical cytoskeletal networks are shown *en face* and all other images are cross-sections. (E,F) Quantification of cortical actin cable density (E) and cortical actin cable length (F) during s10B in genotypes and conditions as shown. E,F share a key. Two-way ANOVA with Tukey’s multiple comparison test. Conditions sharing letters are not significantly different from one another, conditions with different letters are significantly different, p<0.0001, n=15-45 per genotype. (G,H) Quantification of cytoplasmic (G, throughout s10B) and cortical (H, s10B.3) MT density in genotypes and conditions as shown. (G) Two-way ANOVA with Tukey’s multiple comparison test. Conditions sharing letters are not significantly different from one another, conditions with different letters are significantly different, p<0.0001, n=15-45 per genotype (G), p<0.0005, n=5-12 per genotype (H).

Like CLIP and EB1 KDs (Fig. 2B,C), in LatA-treated cells we often observed free cables and coaligning MTs concentrated around the nucleus (Fig. 6B, yellow arrowheads). Acute LatA treatment significantly depleted both the cortical and cytoplasmic MT networks (Fig. 6B, G, H), similar to what we observed when we reduced any one of the three actin bundlers (Chou et al., 2026, Table 1). Because reduction of any one of the filament bundlers did not produce free cables (Fig. 4D-G), it seems unlikely that the loss of MTs is causing actin cable detachment in LatA-treated cells. Instead, those free cables are likely a direct consequence of the strong block to actin filament assembly.

Next, we knocked down LIMK, a negative regulator of Cofilin (Arber et al., 1998; Yang et al., 1998), to enhance Cofilin activity and favor disassembly directly without perturbing assembly. LIMK KD produced significant numbers of free cables (Fig. 5B,F, Fig. 6C, arrowheads) that shrank at a comparable rate to those in Lat-A treated nurse cells (0.03 µm/min; Fig. 5B,G). This suggests that enhanced filament disassembly disrupts the cortical continuity of actin cables and that free cable shrinkage is mediated by Cofilin. Chronically reducing LIMK also significantly reduced cortical cable number and length (Fig. 6E,F), and cortical and cytoplasmic Ace-MT density (Fig. 6G,H). The actin cable and MT defects we observed could be direct and indirect effects of excessive Cofilin activity, but because LIMK is a serine/threonine kinase with multiple potential targets, it is possible that its depletion affects actin cable assembly and MTs independent of Cofilin.

To favor filament assembly, we inhibited disassembly by reducing Cofilin activity through KD of Flare/AIP1 (Flr), a conserved cofactor that enhances Cofilin-mediated actin filament disassembly (Rodal et al., 1999; Goode et al., 2023). Knockdown of Cofilin itself blocked oogenesis prior to s10B. Flr KD significantly increased the fraction of free cables (Fig. 5C,F, Fig. 6D, arrowheads) and virtually all of those appeared to be the result of detachment rather than breakage (Fig. 3J). Because loss of Flr should reduce disassembly and thereby favor assembly, we predicted that the free cables would grow. Remarkably, this is exactly what we saw: free cables in the Flr KD continued to elongate (Fig. 5C,H). Though the rate of elongation was somewhat slower, it was not significantly different from cortically attached Flr KD or control cables (approx. 0.08 µm/min; Fig. 5H, Fig. S3E). Because Dia elongates WT cortical cables, we asked if formins are required for the elongation of free cables by treating Flr KDs with the formin inhibitor SMIFH2 (Rizvi et al., 2009). DMSO treatment alone did not significantly affect the free cable elongation rate in Flr KD nurse cells (Fig. 5D,H), but addition of the formin inhibitor significantly reduced elongation and the free cables maintained stable length (Fig. 5E,H). Together, these data suggest that free cable disassembly is mediated by Flr and Cofilin, while assembly is mediated by formins. Dia is enriched at the nurse cell cortex (Logan et al., 2022; Chou et al., 2026), but we did not observe endogenous Dia to localize along cortical cables or to free cables. However, Drosophila have 6 formins, all of which appear to be expressed in the female germline based on scRNA-seq data (Flybase), suggesting that other formins could be contributing to free cable elongation.

Given the robust growth of the free cables in live Flr KD nurse cells, we were very surprised to find that in fixed cells the free cables were typically short (Fig. 6D, arrowheads) and comparable to what we observed in cells with shrinking free cables (Fig. 2B-D). We previously showed that nurse cell cables elongating from the region of the cortex adjacent to the overlying follicle cells (“follicle cables”) can only be visualized in live cells (Logan et al., 2022). The elongating portion of the free cables in the Flr KD may be similar to wild type follicle cables in some way, rendering them resistant to fixation. In addition to the dramatic increase in free cables, we found that the loss of Flr significantly decreased cortical cable density and length (Fig. 6D-F). The density of the cytoplasmic and cortical Ace-MT networks was unaffected (Fig. 6D,G,H), arguing that the actin cable defects are a result of loss of Flr’s actin disassembly activity rather than a consequence of MT network changes.

Overall, these findings support a model in which the cortical continuity and continued elongation of actin cables are maintained through coordinated Dia- and Ena-mediated assembly and Cofilin- and Flr-mediated disassembly. Disrupting normal assembly-disassembly dynamics by blocking assembly (LatA treatment), or increasing or decreasing disassembly (LIMK KD and Flr KD respectively) leads to a loss of the cortical continuity of actin cables. Likewise, free cables in the cytoplasm appear to be maintained by coordinated formin-mediated assembly and Cofilin- and Flr-mediated disassembly. Here, inhibiting disassembly (Flr KD) produced growing free cables, while promoting disassembly (LIMK KD) or inhibiting assembly (LatA treatment) produced shrinking free cables.

### Free cables and cortical cables exhibit barbed-end disassembly

Typically, the barbed-end of an actin filament favors assembly while the pointed end favors disassembly. Treadmilling describes filament remodeling via the well-known, balanced barbed-end assembly and pointed-end disassembly that maintains the length of a filament while turning over its component subunits (Bugyi and Carlier, 2010). In microvilli, the persistent core bundles of actin filaments exhibit treadmilling with assembly at the barbed ends in the distal tip of the bundle near the plasma membrane and disassembly at pointed ends at the proximal end of the bundle in the cytoplasm of the cell body (Tyska and Mooseker, 2002; Meenderink et al., 2019). Recent studies have shown that actin filaments can disassemble from barbed ends both in vitro and in cells (Romet-Lemonne and Jegou, 2021; Courtemanche and Henty-Ridilla, 2024) driven by regulators including Cofilin, Twinfilin and Srv2/CAP (Wioland et al., 2017; Arya et al. 2024; Johnston et al, 2015, Shekhar et al. 2021; Hakala et al., 2021; Ailmov et al., 2023). The persistent filament bundles in stereocilia exhibit balanced assembly and ADF-AIP1-mediated disassembly at the barbed-ends of the filaments at the distal tip of the filament bundle near the plasma membrane (Drummond et al. 2015; Narayanan et al., 2015). To our knowledge, such barbed-end filament remodeling has not been described in other filaments or filament bundles in vivo, and has not been reported in the context of growing filaments.

To distinguish between treadmilling and barbed-end filament remodeling in the actin cables, we took advantage of shrinking free cables to ask whether they disassemble from the barbed ends, the pointed ends, or both. We blocked the barbed ends of shrinking free cables by treating CLIP KD nurse cells with the barbed-end binding molecule Cytochalasin D (CytoD). We reasoned that if the filaments within the free cables in the CLIP-190 KD nurse cells are normally treadmilling (Fig. 7A), blocking the barbed ends of free cables that normally shrink would leave the pointed ends free to disassemble, and the free cables would continue to shrink. Alternatively, if the free cables are normally undergoing barbed-end filament remodeling (Fig. 7B), blocking the barbed end with CytoD would prevent monomer disassembly, and the formerly shrinking cables would maintain a stable length. In DMSO treated CLIP-190 KD controls, the shrinking free cables continued to shrink (Fig. 7C,G). At 2.5 µg/ml CytoD, we found that the formerly shrinking CLIP-190 KD cables instead maintained a stable length (Fig. 7D,G; the line does not significantly deviate from zero). This suggests that disassembly of the free cables is occurring at the barbed ends of the filaments. Treating wild-type nurse cells with CytoD produced free cables and these also maintained a stable length (Fig. 7E, F, H) consistent with a block to both barbed-end assembly and disassembly.

**Figure 7:**
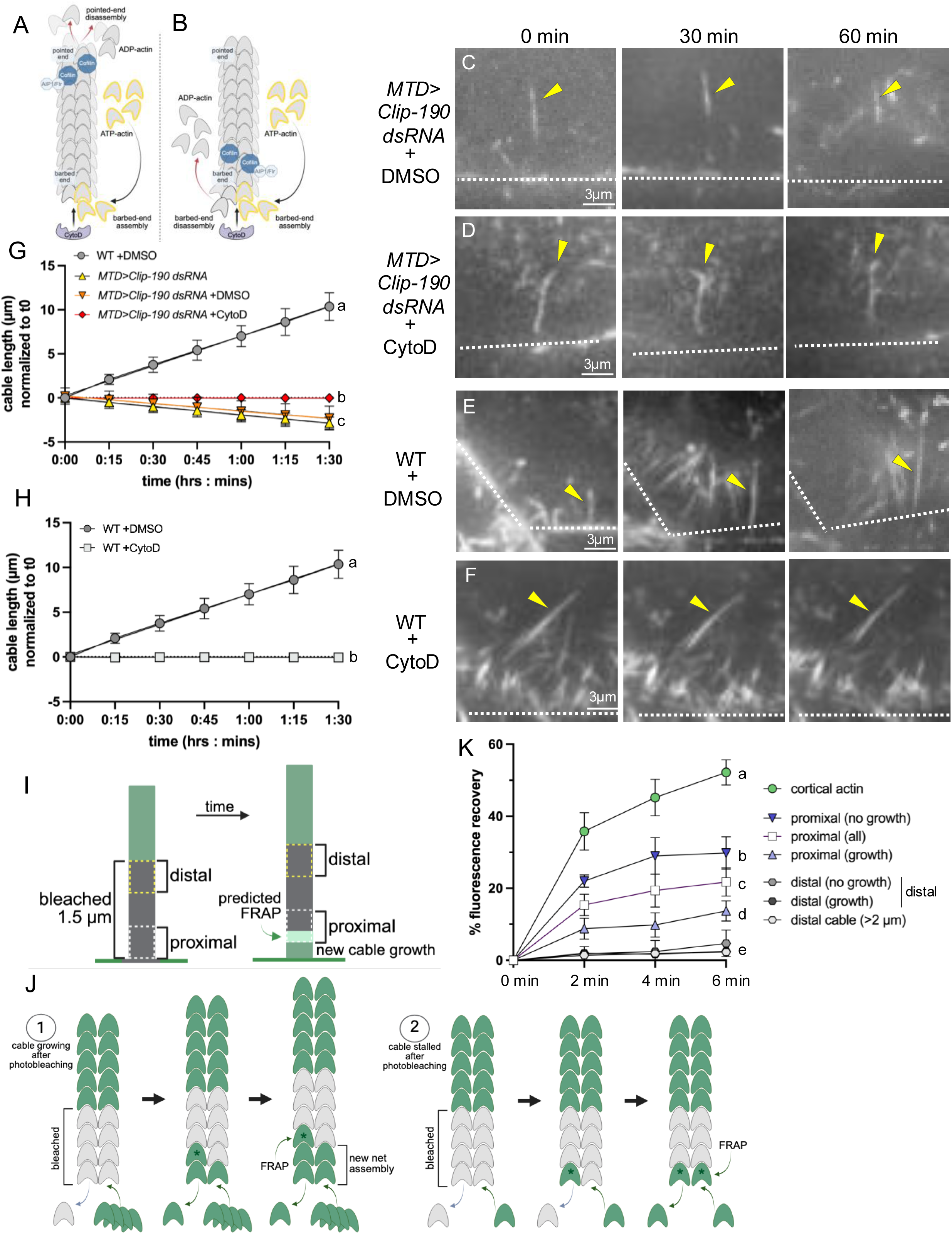
Assessing barbed-end dynamics in free and cortical actin cables. (A,B) Schematic of how Cytochalasin D (CytoD) was used to distinguish between two modes of filament dynamics in shrinking free cables. (A) If free cable actin filaments undergo treadmilling, blocking the barbed end would not prevent pointed-end disassembly, and cables would continue to shrink. (B) Alternatively, if barbed-end disassembly occurs, barbed-end blockade would halt monomer loss, stabilizing cable length. (C-F) Images of s10B actin cables in live nurse cells labeled with SPY-actin. Actin cables were imaged for 90 minutes; representative time points at t=0, t=30, and t=60 minutes are shown. The cortex is indicated with white dashed lines, and a representative cable is indicated by yellow arrowheads. (scale bar = 3 µm) (G,H) Quantification of cable length over time. Initial cable lengths were normalized to 0 µm at time = 0:00 for comparison. In WT, elongating cortical cables were measured. In KDs and drug-treated conditions, cables detach at time zero and free cable lengths were measured. Simple linear regression with ANOVA. Genotypes sharing letters are not statistically different from one another. p<0.0001. n=20∼25 cables per genotype/condition. (I) Schematic diagrams illustrating the region of the cable bleached and the proximal and distal positions where recovery was assessed. ≥1.5 µm of an GFP-Act5C labeled actin cable adjacent to the cortex was photobleached and recovery in a proximal region (0.5 µm) and a distal region (0.5 µm) of the bleached area was assessed over time as shown. The proximal and distal regions were shifted along the cable in concert with new cable growth. The region of the cable that we predict will exhibit fluorescence recovery (predicted FRAP) is shown. (J) Schematic diagrams illustrating the predicted FRAP if cables continue to grow after photobleaching (1), and if cable growth is stalled after photobleaching (2). Asterisks distinguish monomers added during turnover (*) from those added during new assembly. (K) Quantification of FRAP of cortical actin and different regions of WT cortical cables as shown. The “proximal all” condition includes recovery of all proximal cable regions regardless of continued cable growth. Simple linear regression with ANOVA. Conditions with different letters are significantly different from each other. p<0.0001, n=22 cables in total. All schematics created in Biorender.

To visualize barbed-end disassembly in wild type, cortically associated cables, we performed fluorescence recovery after photobleaching (FRAP) on actin cables in s10B nurse cells labeled with GFP-Actin5C (Huelsmann et al., 2013). For all experiments, we set the post-bleach fluorescence intensity to zero and assessed recovery over 6 min. As a control, we showed that nurse cell cortical GFP-Actin5C fluorescence recovered quickly (Fig. 7K, Fig. S4A), consistent with previous reports in other cell types (Fritzsche et al., 2013; Kelkar et al., 2020).

Next, we focused on the proximal region of the actin cables closest to the cortex which corresponds to the barbed ends of the assembling unit bundle (Fig 7I). If no barbed-end disassembly occurs, that bleached region would not show any recovery and over time it would move away from the cortex as the cable elongates. If barbed-end disassembly occurs, we predicted that the proximal bleached region would recover some fluorescence as some bleached subunits are turned over and new fluorescent subunits are added (Fig. 7I). We bleached a 1-2 µm region of the actin cable including the proximal cable end and assessed recovery in a 0.5 µm window immediately adjacent to the cortex (Fig. 7I, proximal). To account for new growth, the recovery window was shifted distally based on the amount of cable growth we observed over the recovery period (Fig. 7I). Net wild type cable growth rate is 0.12 µm/min (approx. 44 monomers/min; Fig. 3H), consistent with our previous work (Logan et al. 2022). Using the shrinkage rate of free cables induced by LatA treatment, we estimated the disassembly rate may be roughly three times slower (approx. 0.04 µm/min or 15 monomers/min; Fig. 5G). Therefore, we predicted that the amount of recovery in the proximal cable would likely be relatively modest. As an internal control, we also assessed recovery in a 0.5 µm distal window within the same bleached region (Fig. 7I, distal). Our model predicts that monomers in more distal regions of the cable do not turnover and thus we would not see any fluorescence recovery. Consistent with our predictions, we observed significant fluorescence recovery in the proximal regions of the bleached cables (Fig. 7K, proximal–all) and virtually no recovery in the matched distal regions of the same cables (Fig. 7K, distal). As an additional control, we bleached regions at least 2 µm distal to the cortex and found that these unmatched distal bleached regions likewise did not exhibit detectable fluorescence recovery (Fig. 7K, distal cable, Fig. S4B; Huelsmann et al. 2013).

Because cable elongation is relatively slow, we typically use 15 min intervals for live imaging to minimize tissue damage. Based on the 0.12 µm/min growth rate we calculated from those experiments, we predicted that the bleached cables would grow approx. 0.7 µm in the 6 min we assessed fluorescence recovery. Half of the bleached cables elongated 0.5–1 µm post-bleach while the other half appeared to be stalled with no significant elongation during that time. Our data suggest that if the filaments in those cables were experiencing no assembly, the cables would detach from the cortex as they do in LatA treated cells (FIg. 5F). Thus, we reasoned that those stalled cables could be experiencing balanced, equivalent barbed-end assembly and disassembly and therefore no detectable elongation. If that is the case, we predicted that the proximal bleached regions in stalled cables would have more opportunity to turnover the bleached subunits resulting in significantly higher fluorescence recovery (Fig. 7J, compare 1 and 2). That is what we observed (Fig. 7K, proximal–growth vs proximal–no growth). The recovery of the matched distal regions in the two groups was not significantly different (Fig. 7K, distal–growth vs distal–no growth). Additional imaging with higher temporal resolution is needed to determine whether stalling is a normal cable growth phenomenon. Taken together, the results of our CytoD and FRAP experiments suggest that the actin filaments within both shrinking free cables and elongating wild-type cortical cables undergo barbed-end disassembly.

### Capping Protein protects actin cables from disassembly

For filaments to elongate and cables to grow when both assembly and disassembly are occurring at the cortical barbed ends of the filaments, the barbed end must be tightly regulated such that the rate of filament assembly exceeds the rate of disassembly. One regulator of the barbed end is Capping Protein (CP). In vitro studies demonstrated that CP antagonizes formin at the barbed end through the formation of a CP-formin “decision complex” (Bombardier et al., 2015). Consistent with a role in limiting assembly, loss of CP can lead to: excessive filopodia formation (Mejillano et al., 2004; Gates et al., 2007), elevated levels of actin in Drosophila imaginal epithelia (Delalle et al., 2005; Janody & Treisman, 2006) and defects in bristle morphogenesis are consistent with excessive actin assembly (Hopmann et al., 1996). In yeast, loss of CP resulted in the expansion of actin patches (Wirshing et al., 2023). CP could also protect the barbed end from disassembly (Caldwell, 1989; Wear et al. 2003), but in vivo there is limited evidence for the loss of actin structures in the absence of CP. Interestingly, loss of CAPZb in hair cells increases actin density at the adherens junctions but also results in short, thin, or absent stereocilia (Avenarius et al., 2017). This suggests a role for CP in protecting actin filaments from disassembly in some actin bundles.

Based on these findings, we asked whether CP regulates filament barbed-ends in growing actin cables. Prior to s9, CP is diffusely cytoplasmic in nurse cells without any apparent subcellular enrichment (Fig. 8A,D). Starting in s9, we detected cortical enrichment that increased through s10A and plateaued at s10B (Fig. 8B-D). Interestingly, Ena exhibited a similar increase in cortical enrichment during this same developmental window (Fig. 8A-C,E). Because the cables are composed of bundles of 2-4 µm unit bundles (Guild et al. 1997), we considered the possibility that CP could be found at the barbed ends of the unit bundles periodically along the cables. However, we did not detect CP accumulation along the cables (Fig 8C and Fig. S5A). These results suggest that CP may be functioning primarily in the actin cortex and at the cortical barbed ends of the elongating actin cables.

**Figure 8:**
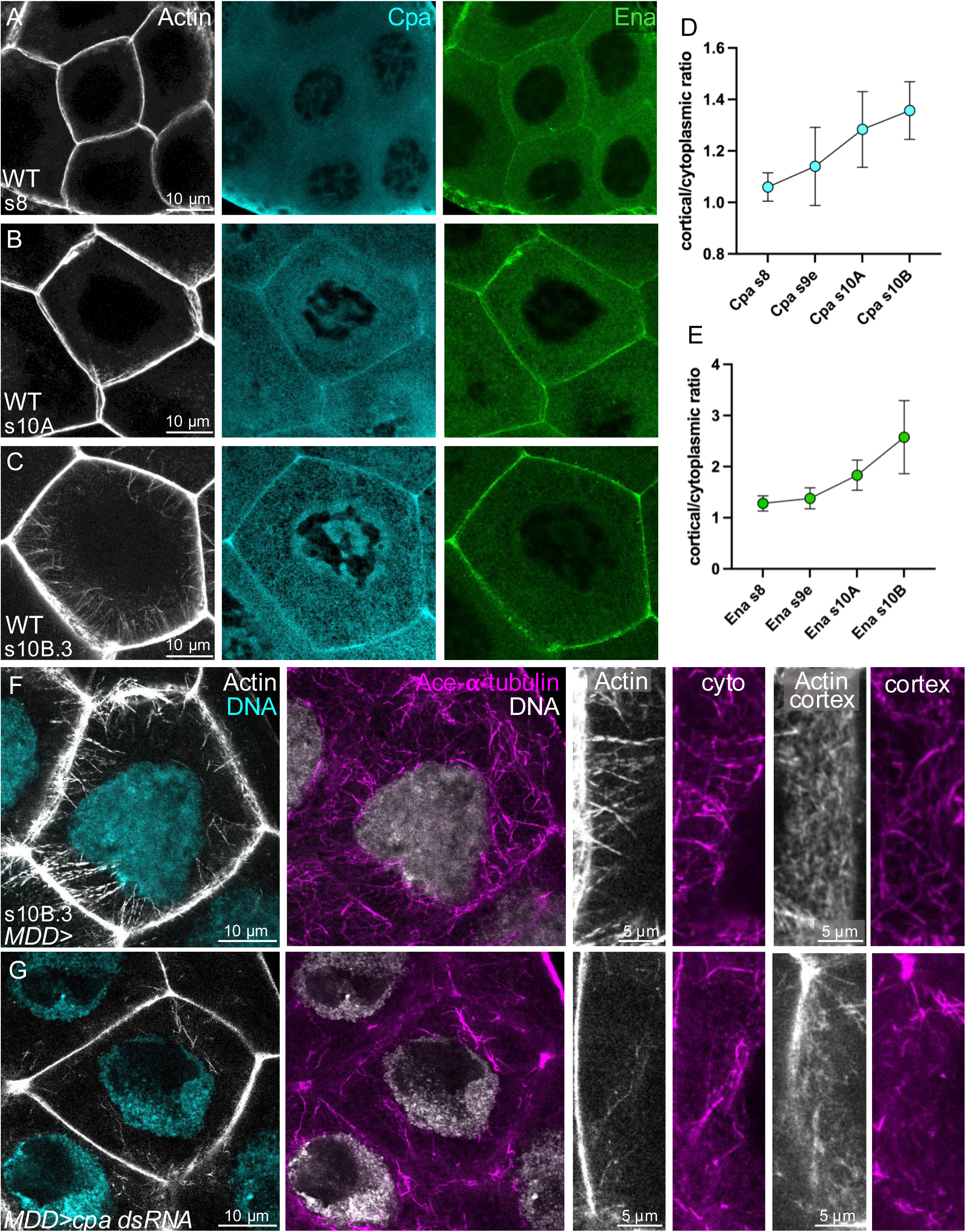
Capping protein (CP) becomes cortically enriched at s10B and is required for cortical actin cable assembly. (A-C) Immunofluorescent images (max projection of three 0.4 µm z-slices) of WT nurse cells in cross section at s8 (A), s10A (B) and s10B.3 (C) triple-labeled for Actin (grey; phalloidin), Capping Protein (Cpa, cyan) and Ena (green) (scale bar = 10 µm). (D,E) Quantification of the cortical to cytoplasmic ratio of fluorescent intensity for Cpa (D) and Ena (E) at the indicated stages. (F,G) Representative images of fixed s10B.3 nurse cells triple-labeled for actin cables (grey; phallodin), DNA (DAPI; grey or cyan) and the Ace-MT network (magenta; anti-Ace-α-tubulin) in WT (F), and *MDD>cpa dsRNA* (G) nurse cells. These are shown at lower magnification (max projection of two 0.4 µm z-slices; scale bar = 10 µm) and higher magnification (max projection of seven 0.4 µm z-slices; scale bar = 5 µm). All images were taken from the same nurse cells. The cortical cytoskeletal networks are shown *en face* and all other images are cross-sections.

Previous work by Gates et al. (2009) showed that *cpb* mutant nurse cells had disorganized actin cables. We found that depleting CP via KD of either one of its subunits (Cpa or Cpb) significantly reduced total cable numbers and their length, suggesting reduced cable initiation and slower filament elongation (Fig. 6E,F, Fig. 8F,G, Fig. S1K), similar to what we have observed in KDs of Dia, Ena or the actin filament bundlers (Chou et al., 2026). Loss of CP also significantly increased the proportion of free cables (Fig. 5F). Consistent with a role for CP in protection against disassembly, we found that free cables shrank in CP KDs (Fig. S5E,H), and this was rescued by acute CytoD treatment blocking the barbed ends (Fig. S5F-H). All these data are consistent with the model that CP is functioning at the cortical barbed ends of actin filaments within the growing actin cables. CP may regulate both barbed-end assembly and disassembly, but the net effect appears to be protection of those ends from excessive disassembly.

Because loss of CP caused a dramatic change in the appearance of cortical actin suggesting changes in cortical filament organization (Fig. 8F,G, Fig. S5C,D), it was not surprising that cortical MT density significantly decreased (Fig. 6H, Fig. S5B). In cultured cells, CAPZ reduction increased the thickness of cortical actin with an associated decrease in cortical tension (Chugh et al., 2017). The density of the cytoplasmic MT network was also significantly reduced in CP KDs (Fig. 6G). This reduction in both cortical and cytoplasmic MTs is similar to what we observed when we reduced any of the three actin filament bundlers (Quail/Villin, Singed/Fascin, or Fimbrin; Chou at el., 2026). While the bundler KD cells also had a strong reduction in cable number (Chou et al., 2026) similar to CP KDs (Fig. 6E), they had significantly fewer free cables than control cells (Fig. 4G). This suggests that the free cables in CP KD cells are unlikely to be the result of the loss of MTs.

## Discussion

### How to build actin cables: Actin filament assembly and disassembly are required for persistent cable elongation and cortical continuity

Here we have shown that the persistent elongation of actin cables and their association with the cell cortex is dependent on the availability of actin monomers for filament assembly and, surprisingly, on mechanisms that regulate the disassembly of actin filaments. Blocking assembly or blocking or enhancing disassembly all caused actin cables to detach from the cortex. This and the subsequent behavior of the free cables in the cytoplasm has yielded insights into the mechanisms underlying cable assembly. Our results suggest that the assembly and disassembly required for actin cable elongation are occurring at the cortical barbed-ends. These barbed-end dynamics are necessarily imbalanced– in order for cables to elongate, the monomer on-rate must exceed the off-rate (Fig. 9). Dissecting the mechanisms regulating and coordinating assembly/disassembly is complicated by multiple modes of crosstalk between the actin and MT cytoskeletons. The effects of all our genetic and inhibitor experiments on actin cables and the MT network are summarized in Table 2.

**Figure 9:**
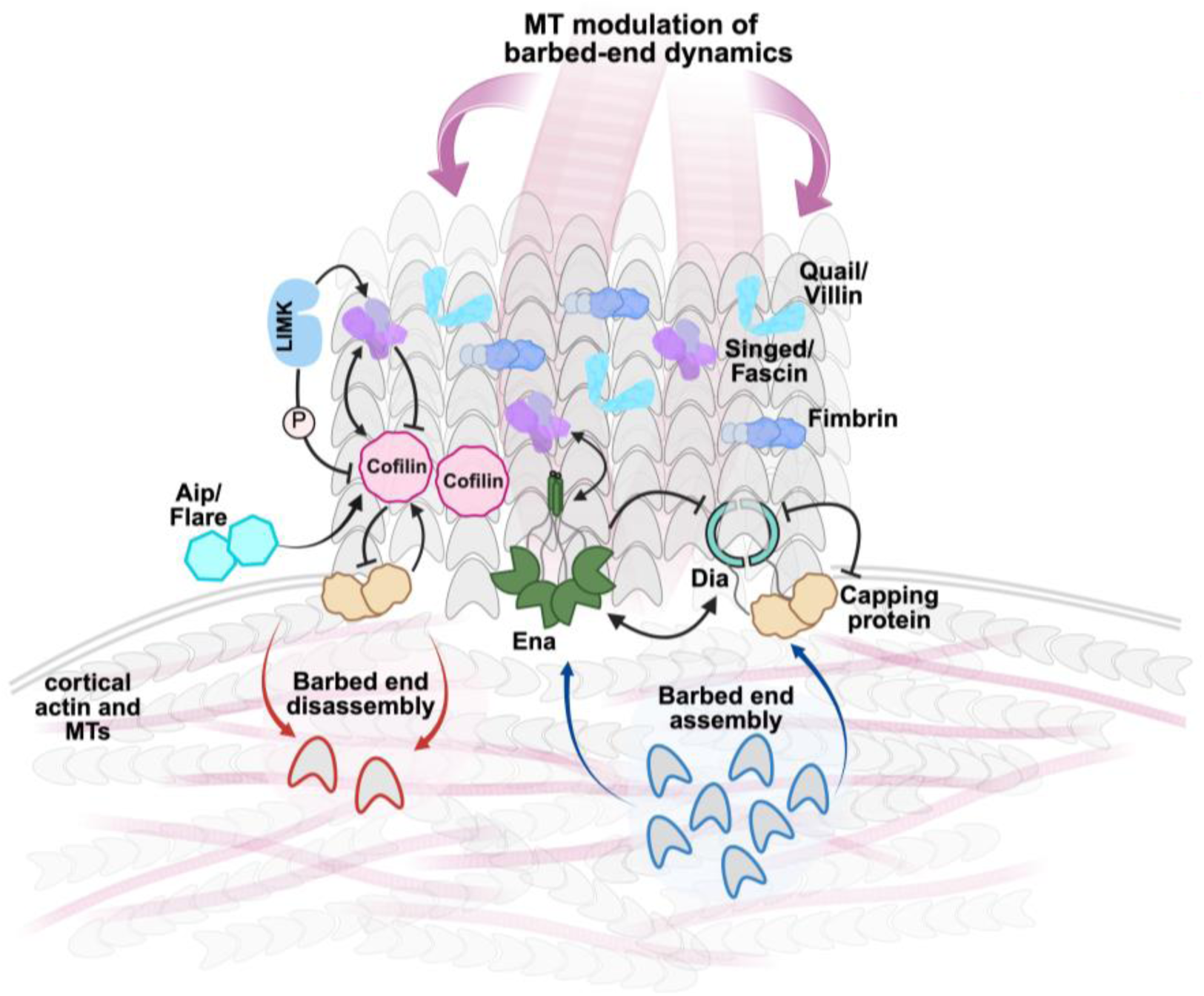
Barbed-end assembly and disassembly modulated by MTs are required to elongate cortical actin cables. Our results suggest that both barbed-end assembly and disassembly are required for proper actin cable formation and elongation. We propose that barbed-end disassembly serves to regulate filament length, ensuring uniformity among the filaments bundled by Singed/Fascin, Quail/Villin and Fimbrin. This uniformity may be critical for forming well-bundled, structurally coherent actin cables. Our data suggest that the disassembly factor Cofilin and its regulators Aip/Flare and LIMK are necessary for persistent cortical cable elongation mediated by Dia and Ena. Capping Protein (CP) is required to maintain cable elongation and may coordinate barbed-end dynamics contributing to the imbalance between assembly and disassembly. MTs appear to regulate barbed-end dynamics through multiple mechanisms (see Discussion). Direct or indirect activating/coordinating interactions illustrated: Aip/Flare and Cofilin (Rodal et al., 1999), Ena and Fascin (Harker et al., 2019), Cofilin and Fascin (Breitsprecher et al., 2011), Cofilin and CP (Wioland et al., 2017) and LIMK and Fascin (Jayo et al., 2012). Direct or indirect inhibitory interactions illustrated: LIMK and Cofilin (Arber et al., 1998; Yang et al., 1998), Cofilin and CP (Wioland et al., 2017), Dia and CP (Bombardier et al., 2015), and Fascin and Cofilin (Chikireddy et al., 2024). Previous work demonstrated an inhibitory interaction between Ena and Dia (Bilancia et al., 2014), but our work suggested a collaborative interaction in actin cable assembly (Logan et al., 2022). Created in Biorender.

**Table 2:** Summary of the effects of all of genetic and inhibitor experiments on actin cables and the MT network. Asterisks indicate data that were previously published in Chou et al., 2026. “Manipulation” indicates the known or predicted effects of the genotype and/or inhibitor treatment.

The most direct ways that we manipulated the filament barbed-ends or biased the system toward assembly or disassembly were through Flr KD, and LatA or CytoD treatment. With Flr KD and consequent reduction of Cofilin activity, we expect that disassembly is impaired while assembly is unaffected. This loss of disassembly produced free cables that underwent formin-dependent elongation in the cytoplasm (Fig. 5H) suggesting that disassembly may be necessary to maintain persistent cortical cable assembly and elongation. Consistent with that notion, the remaining cortical cables were significantly shorter in Flr KD cells (Fig. 6F) reflecting a slightly slower elongation rate (Fig. 5H). Interestingly, this was the only manipulation of actin we performed that did not reduce the density of the MT network (Fig. 6G,H).

By binding to G-actin, LatA reduces filament assembly thereby favoring disassembly. We propose that LatA treatment causes free cables because the assembly rate drops below a critical threshold releasing the cables from the processive filament elongation by cortical Dia and Ena. Although Dia or Ena KD significantly reduced the cable assembly rate (Logan et al., 2022), cable continuity and elongation continued (Fig. 4). We predict that the LatA-induced free cables shrink because the disassembly machinery, including Cofilin and Flr, continues to operate at those barbed ends. Like LatA, treatment with CytoD blocks barbed-end assembly and releases cables from the cortex (Fig. 7F,H). These free cables maintain a stable length in the cytoplasm because CytoD also protects the barbed end from disassembly. We did not assess the MT network in cells treated with CytoD, but in cells treated with LatA the MT network was strongly depleted (Fig. 6G,H). This raises the possibility that the free cables in LatA-treated cells are an indirect effect of the loss of assembly and instead a direct consequence of MT disruption. While we can’t resolve this conundrum, we do know that similar levels of MT depletion in cells with reduced actin filament assembly and bundling do not produce free cables (Chou et al, 2026; Table 1). Thus, we favor the interpretation that free cables in LatA-treated cells are the result of strongly reduced barbed end monomer addition.

The requirement for filament disassembly in the assembly and elongation of actin cables was unexpected. One possible explanation for this apparent contradiction was inspired by in vitro work examining the coordination of filament elongation by Ena and filament bundling by Fascin (Winkelman et al., 2014; Harker et al., 2019; Fig. 9): Ena’s processivity increased with the number of Fascin-bundled filaments, and processivity was enhanced ∼3-fold on shorter, trailing filaments. This provides a potential mechanism to coordinate filament lengths within bundles. If it is important that the lengths of the filaments within a bundle are uniform to some degree, then filament disassembly at the barbed end is an additional way to accomplish this (Fig. 9). Barbed-end disassembly has been observed in vitro and in vivo (Romet-Lemonne and Jegou, 2021; Courtemanche and Henty-Ridilla, 2024), but appears to be less common than pointed-end disassembly. The results of our CytoD and FRAP experiments support the hypothesis that disassembly is occurring at the barbed ends (Fig. 7). We propose that barbed-end disassembly may function as a form of filament sculpting, trimming overly long filaments to generate more uniform lengths that are conducive to tight, organized bundles (Fig. 9). Thus, loss of regulated disassembly at the barbed end could lead to poorly structured cables, interruption in persistent and coordinated filament assembly and bundling, and ultimately, detachment from the cortex.

Barbed-end disassembly must be tightly controlled to enable cable growth. When we increased Cofilin activity by reducing LIMK, cables detached and shrank (Fig. 5G). Cofilin is typically thought of as promoting filament disassembly at the pointed end, but it can promote filament depolymerization at both ends in vitro (Wioland et al 2017, 2019). When filaments are capped by CP, ADF/Cofilin can decorate filaments up to the barbed end, displacing CP resulting in barbed end depolymerization (Wioland et al., 2017; Fig. 9). Our data are consistent with a role for CP in restricting barbed-end disassembly during cable growth. While in most contexts, loss of CP resulted in excessive actin assembly (e.g. Gates et al., 2007; Janody & Treisman, 2005; Wirshing et al., 2023), CP KD in nurse cells reduced cable numbers and their elongation and produced shrinking free cables (Fig. 5G, Fig. 6E,F). Although reduced actin assembly and enhanced disassembly appear to be the net effects of CP loss, it seems likely that CP could play a key role in maintaining the regulated imbalance between assembly and disassembly that enables cable elongation. The depolymerase Twinfilin is another candidate for coordinating barbed-end dynamics. Recent work demonstrated that CP, mDia1 and Twinfilin can simultaneously bind filament barbed ends in vitro (Ulrichs et al., 2023; Reddy et al., 2025), and Twinfilin can displace CP and promote formin-mediated assembly (Ulrichs et al. 2023). Further, Twinfilin promotes Coflin’s binding to the sides of filaments and enhances Cofilin-mediated severing in vitro (Arya et al. 2025; Shekhar et al. 2021). In vivo, Twinfilin localizes to regions of active actin remodeling that are enriched in barbed ends, such as the tips of stereocilia (Rzadzinska et al., 2009), lamellipodia (Hakala et al., 2019), cortical actin patches in yeast (Goode et al., 1998), and ends of actin bundles in developing Drosophila bristles (Wahlström et al., 2001). Notably, Twinfilin appears to be enriched at the cortex of s10B nurse cells (Wahlström et al., 2001), consistent with our model for barbed-end disassembly of actin cable filaments at the cortex.

### Microtubules may promote both filament assembly and disassembly to maintain the cortical continuity of growing actin cables

In the hours preceding the onset of actin cable assembly, the MT network undergoes several dramatic changes (Chou et al., 2026). An anterior to posterior wave of acetylation starting at s9 and ending at the onset of s10B leaves the MT network completely acetylated. This is coincident with a precipitous drop in plus end growth and enhanced MT stability. MTs are required to promote actin cable initiation and elongation (Chou et al., 2026), suggesting that they stimulate actin filament assembly. Here we show that many of the cables that initiate when MTs are reduced detach from the cortex. Every condition affecting MTs directly produced free cables, but surprisingly some conditions produced shrinking free cables (CLIP or EB1 KD) while others produced free cables that maintained a stable length (γ-tubulin KD, nocodazole treated or taxol treated). Overall, our data suggest that MTs may be impacting actin filament dynamics in different, and potentially opposing ways.

We found that both EB1 and CLIP are required for the assembly and/or maintenance of the s10B MT network (Fig. 2). Because MTs are required for actin cable assembly, it was not surprising that actin cable numbers were reduced and that cable elongation was curtailed in EB1 and CLIP KDs. The shrinking free cables we observed in CLIP and EB1 KD appeared similar to the LatA (reduced assembly) and LIMK (enhanced disassembly) phenotypes. Shrinking free cables suggest that the normal imbalance between assembly and disassembly now favors disassembly, and that can be achieved by reducing assembly or enhancing disassembly.

Because loss of CLIP or EB1 strongly reduced the density of the MT network, we predicted that comparable reductions to the network achieved through other perturbations would also produce shrinking free cables. Thus, we were very surprised to find that the free cables produced by γ-tubulin KD or nocodazole treatment maintained a stable length in the cytoplasm, a phenotype similar to that observed following treatment with CytoD. In the case of CytoD, the stability likely results from a block to all barbed-end dynamics. Alternatively, stable free cable length could be the result of balanced assembly and disassembly. In the FRAP experiment, we observed a population of stalled, wild-type cables that maintained their cortical association, at least for the short 6 min duration of the experiment. If that stalling reflects balanced assembly and disassembly, then stable, free cables are more likely to be the result of the cessation of barbed-end dynamics. If this is the case, it argues that MTs can stimulate both assembly and disassembly.

Why free cable fate differs depending on the means of reducing the MT network is not clear. It is possible that the MTs remaining under these different conditions could have significant differences in MT-associated proteins or other characteristics that impact the actin cables. The fact that free cable-MT coalignment was unchanged in CLIP and EB1 KDs and significantly increased following γ-tubulin KD (Fig. 2J) is consistent with the notion that the remaining MTs under these different conditions may have distinct properties. Remarkably, stabilization of the MT network with taxol also produced free cables with stable length (Fig. 3I). This was perplexing as we previously demonstrated that the s10B MT network is highly acetylated, has few growing plus ends, and individual MTs persist for >10 min (Chou et al., 2026); all these properties are consistent with a stabilized MT network. However, Cryo-EM studies have revealed that taxol stabilization and acetylation represent mechanistically distinct modes of MT stabilization, each with different effects on MT structure and mechanical properties (Kellog E.H et al., 2017; Eshun-Willison et al., 2019). These differences may underlie the divergent effects of taxol-stabilized vs acetylated MTs on actin cables. Taken together, our results support a model in which MTs modulate barbed end dynamics to promote the initiation of actin cables and to maintain their persistent elongation from the cortex to the nucleus (Fig. 9).

## Methods

### Immunofluorescence

Flies 1-2 days after eclosion were fed wet yeast paste 24∼48 hours to promote egg production. Ovaries were dissected in freshly prepared Grace’s media (Grace’s medium powder (Sigma, G9771; VWR 95037-608) + 10% Fetal Bovine Serum (FBS) + 7.5% Sodium Bicarbonate (pH 6.2∼6.8), fixed for 20 min in 4% EM-grade paraformaldehyde (PFA) diluted in phosphate-buffered saline (PBS) from an 8% PFA Aqueous Solution (Electron Microscopy Sciences, 157-8) and blocked for 1 h in PBS plus 0.2% Triton X-100 and 4% normal goat serum (NGS) at room temperature. Primary antibody incubation was in PBS plus 0.1% Triton X-100 and 1% NGS with 1:500 Alexa488-Phalloidin (Thermo Fisher Scientific, A12379), 1:350 monoclonal acetylated-⍺-tubulin (Sigma, MABT868; Santa Cruz sc-23950) at 4°C overnight, followed by 1:1000 goat anti-mouse Alexa Fluor™ 568 (Thermo Fisher Scientific A11031) for 3 hr and DAPI (1µg/ml) for 30 min before mounting (Prolong Gold, Invitrogen P36934). The same protocol was carried out for immunofluorescence using 1:350 rabbit monoclonal anti-capping protein (gift from Dr. Jörg Großhans, Philipps University of Marburg) with 1:1000 goat anti-rabbit Alexa Fluor™ 568 (Thermo Fisher Scientific A11011), and 1:500 mouse monoclonal anti-Ena (5G2; DSHB) with 1:1000 goat anti-mouse Alexa Fluor™ 647 (Thermo Fisher Scientific A21235).

### Live imaging of actin cables and drug treatments

For live imaging experiments, s10B egg chambers were isolated as described previously (Spracklen and Tootle, 2013). Egg chambers were imaged in live imaging media consisting of Schneider’s media with 20% fetal bovine serum (FBS), 5 μg/ml insulin, 2 mg/ml trehalose, 5 μM methoprene, 1 μg/ml 20-hydroxyecdysone and 50 ng/ml adenosine deaminase in glass-bottom petri dishes (Azer Scientific ES56291). Egg chambers were labeled with 1:500 SPY-actin (Cytoskeleton, CY-SC202) and 1:1000 Verapamil for 30 minutes incubation in 29°C and washed three times with imaging media prior to imaging (Logan et al., 2022).

Drug treatments were added to live imaging media at the start of imaging (time zero) and egg chambers were maintained *ex vivo* in the imaging media +drug. Concentrations were as follows: Latrunculin A: 1µM. Nocodazole: 7.5µg/ml. Taxol: 2µM. SMIFH2: 200µM. Cytochalasin D: 2.5 µg/ml. DMSO was added accordingly.

### Image acquisition

#### For fixed imaging

samples were imaged on a Olympus Fluoview 3000 confocal microscope with a X60/1.42NA. Fixed images were acquired with Fluoview software (FV-10 ASW3.1, Olympus) at 0.4-0.43 µm step size, and a projection of two optical slices was used to show representative images. For Fig. 2B-D, images were acquired with a Prime95B CMOS camera (Photometrics) on a Zeiss Axiovert 200 M microscope with an X-Light V2 spinning disc scan head (Crest Optics). Fluorophores were excited with a Celesta Light Engine (Lumencor). Images were acquired with MetaMorph software at 0.2 µm step size, and a projection of five optical slices was used to show representative images.

#### For live imaging (excluding FRAP)

samples were imaged with an Evolve EMCCD on a Zeiss Axiovert 200 M microscope with an X-Light V2 spinning disc scan head (Crest Optics). Fluorophores were excited with a Celesta Light Engine (Lumencor). Images were acquired with MetaMorph software at 0.5 µm step size.

#### For FRAP assays

samples were imaged on a Olympus Fluoview 3000 confocal microscope with a X60/1.42NA. Images were acquired with Fluoview software (FV-10 ASW3.1, Olympus) at 0.6 µm step size. GFP-Actin was photobleached using 100% laser power of the 488-nm laser for 5 seconds using the tornado bleach tool.

### Image Analysis

All experiments were replicated 2-3 times. n refers to either number of egg chambers or cytoskeletal components as specified. Basic image processing and measurements were performed with Fiji (Image J v2.14; National Institutes of Health) and Adobe Photoshop (v25.9.1). Prism 10 (GraphPad) was used to generate all graphs and perform statistical analyses. Microsoft Powerpoint was used to prepare all figures. Schematics and models were created with BioRender.

Methods for quantifying actin cable density and length, cortical and cytoplasmic Ace-MT density, actin cable and Ace-MT coalignment, and cortical fluorescence intensity were previously described (Chou et al., 2026).

### FRAP

*UASp-GFP-Act5C* was expressed using the *mat-tub-gal4_GeneSwitch* driver without RU486 induction as in Huelsmann et al., 2013. Stronger expression of GFP-Act5C with other maternal drivers created too much background fluorescence and cables could not be resolved.

#### Actin cortex and distal cable fluorescence recovery

A 2-6 µm region of the cortex or a distal region of an actin cable (>2 µm from the cortex) were bleached and recovery was assessed over the entire bleached region. At each time point, the average fluorescence intensity in each recovery window was measured and normalized to the cytoplasmic fluorescence (fluorescence intensity averaged from 5 regions (100-400 µm^2^) of surrounding cytoplasm was averaged).

#### Proximal cable fluorescence recovery

A 1-2 µm region of the actin cable overlapping the cortex was bleached and recovery was assessed in two regions: “proximal”– a 0.5 µm window immediately adjacent to the cortex, and “distal”– a 0.5 µm window starting 0.5 µm distal to the proximal window (Fig. 7I). If the cable continued to elongate during the 6 min recovery period, the proximal and distal recovery windows were shifted distally based on the amount of cable growth observed (Fig. 7I). At each time point, the average fluorescence intensity in each recovery window was measured and adjusted to the cytoplasmic fluorescence (fluorescence intensity in 5 regions (x µm) of surrounding cytoplasm was averaged).

### Plasmid constructs

CLIP-190 dsRNA (1) targeting sequence GGAGCCACTGCCTCCAAATTA CLIP-190 dsRNA 2) targeting sequence CGTGAAAGCGATGACAATTTA. The oligos of CLIP-190-shRNA (1) (tagcagt - GGAGCCACTGCCTCCAAATTA - tagttatattcaagcata - TAATTTGGAGGCAGTGGCTCC -gcg), and CLIP-190-shRNA (2) (ctagcagt - CGTGAAAGCGATGACAATTTA - tagttatattcaagcata - TAAATTGTCATCGCTTTCACG -gcg) were inserted into the pWalium22 vector (pWALIUM22 (DGRC Stock 1473; https://dgrc.bio.indiana.edu//stock/1473; RRID:DGRC_1473)). Constructs were sent to BestGene for injection via phiC31 integrase-mediated transformation (CLIP190-dsRNA (1) at attP40 docking site; CLIP190-dsRNA (2) at attP2 docking site).

### Semi-quantitative reverse transcriptase PCR

Whole ovaries were dissected in Grace’s insect medium as described above and homogenized in TRIzol reagent (Zymo Research). Total RNA was isolated according to the manufacturer’s instructions, followed by RNase-free DNase treatment overnight. cDNA synthesis was conducted using the SuperScript™ IV First-Strand Synthesis System (Thermo Fisher Scientific). RT-PCR was performed using primers specific for *CLIP-190*, *Eb1*, *LIMK*, and *flr*. *CLIP-190* (forward: 5’-TACGCTATTTGGGAGAGACACA-3’; reverse: 5’-TTTTTGCCGCTAGGCTCATCC-3’). *Eb1* (forward: 5’-ACCTGGAGGGATTGGAAAAGG-3’; reverse: 5’-GTCATCGGCTTCTTGGCAAA-3’). *LIMK* (forward: 5’-GTGAACGGCACACCAGTTAGT-3’; reverse: 5’-ACTTGCACCGGATCATGCTC-3’). *flr* (forward: 5’- AGCACTCGTGTGCAGTGAAC-3’; reverse: 5’-GTGTCCCAAATGCGAATCTT-3’). Reference gene Ribosomal protein L32 (*RpL32*) (forward: 5’-AGCATACAGGCCCAAGATCG-3’; reverse: 5’-TGTTGTCGATACCCTTGGGC-3’). Changes in transcript levels were evaluated by quantifying PCR band intensities on agarose gels and normalizing them to *RpL32* expression.

### Drosophila genetics

Fly stocks and crosses were maintained on standard cornmeal-molasses food supplemented with dry active yeast. Fly stocks were kept at room temperature. To induce expression using Gal4 drivers, crosses using genotypes marked with asterisks* were kept at 29°C, others not indicated were raised at room temperature.

| Genotype | Source/Identifier | Chromosome information |
|---|---|---|
| *Oregon-R* (wildtype control) | BDSC |  |
| *MDD-GAL4 (maternal double driver)* | BDSC_80361 | *y<sup>1</sup>w;P{matalpha4-GAL-VP16}67; P{matalpha4-GAL-VP16}15* |
| *MTD-GAL4 (maternal triple driver)* | BDSC_31777 | *P{w[+mC]=otu-GAL4::VP16.R}1, w; P{w[+mC]=GAL4-nos.NGT}40; P{w[+mC]=GAL4::VP16-nos.UTR}CG6325[MVD1]* |
| *\*UASp-γtubulin 23C-dsRNA* (TRiP.GL01171) | BDSC_42799 | attP2, III |
| *\*UASp-γtubulin37C-dsRNA* (TRiP.HMS00517) | BDSC_32513 | attP2, III |
| *UASp-CLIP-190 dsRNA (1)* | This study | II |
| *UASp-CLIP-190 dsRNA (2)* | This study | III |
| *UASp-EB1 dsRNA* | BDSC_36599 | attP2, III |
| *UASp-EB1 DN* | BDSC_58722 | {UASp-mCherry.Eb1.DN}ZH-86Fb (III) |
| *\*UASp-dia-dsRNA(1)* (TRiP.HMS00308) | BDSC_33424 | attP2, III |
| *\*UASp-ena-dsRNA(1)* (TRiP.HMS01953) | BDSC_39043 | attP2, III |
| *qua[1]cn[1]bw[1]speck[1]/CyO* | BDSC_3350 | II |
| *qua[6-396]/SM1* | BDSC_4571 | II |
| *In(1)Mud, Mudd[1]/In(1) dl-49 sn[X2]* | BDSC_1035 | X |
| *\*UASp-Fim-dsRNA(1)* (TRiP.HMS00729) | BDSC_32936 | attP2, III |
| *UASp-LIMK-dsRNA (1)* | BDSC_42576 | attP40, II |
| *UASp-LIMK-dsRNA (2)* | BDSC_57012 | attP2, III |
| *UASp-flr-dsRNA (1)* | BDSC_42788 | attP2, III |
| *UASp-flr-dsRNA (2)* | BDSC_31145 | attP2, III |
| *UASp-cpa-dsRNA* | BDSC_31124 | attP2, III |
| *UASp-cpb-dsRNA* | BDSC_50954 | attP40, II |
| *UASp-GFP-Act5C* (Roper et al, 2005) | from S. Huelsmann |  |
| *mat-tub-gal4_GeneSwitch* (Huelmann et al., 2013) | from S. Huelsmann |  |
| *\*UAS-shot<sup>ABD</sup>-RNAi* | Lu et al., 2016 | attP2, III |
| *\*UAS-shot<sup>ROD</sup>-RNAi*<br>(UASp-shot dsRNA TRiP.GL01286) | BDSC_41858 | attP2, III |
| *\*UASp-Patronin-dsRNA* | gift from Dr. Jörg Großhans, Philipps University of Marburg | attP2, III |
| *\*UASt-Patronin-dsRNA* |  | II |
| *matatub-Gal4-VP16[V37], Rhi RNAi* | Dr. Vladimir Gelfand, Northwestern University | III |

## Supplementary Tables and Figures

**Figure S1: Knockdown efficiency and validation of knockdown phenotypes**

(A) Agarose gels displaying the results of RT-PCR performed to assess dsRNA efficacy in *MTD>CLIP-190 dsRNA*, *MTD>Eb1 dsRNA, MTD>LIMK dsRNA,* and *MTD>flr dsRNA* ovaries*. rpl32* is a housekeeping gene.

(B-E) Quantification of total transcript levels from (A). RNA was extracted from whole ovaries. Because knockdown is only occurring in the germline and not in the somatic follicle cells, overall transcript levels may appear higher than expected for the knockdown.

(F) Quantification of the ratio of cortical to cytoplasmic fluorescence intensity (FI) for CP across s10B in control (*MTD>*) and CP knockdown (*MDD>cpa dsRNA*) nurse cells. s10B.1 and s10B.2 were grouped, and s10B3 and s10B.4 were grouped. One-way ANOVA with Tukey’s multiple comparison test, ****p<0.0001. n=3-6 egg chambers per genotype per substage therefore n=6-12 egg chambers per group.

(G-J) Independent dsRNA or dominant-negative transgenes for CLIP, Eb1, and γ-tubulin were expressed to validate representative phenotypes. Cytoplasmic Ace-MT density (G), actin cable density (H), cable length (I) and the proportion of free vs cortical cables (J, combined s10B.1-s10B.4) were quantified. Data for controls, *MTD>CLIP dsRNA (1)*, *MTD>Eb1 dsRNA*, and *MTD>γ-tubulin 23C dsRNA* are also presented in Fig. 2E-G,I. Statistics as described in Fig. 2.

(K,L) Independent dsRNA transgenes targeting LIMK and CP were expressed to validate representative phenotypes in combined s10B.1-s10B.4. Cortical cable density (K) and cytoplasmic MT density (L) were quantified. Data for control, *MTD>LIMK dsRNA*, and *MTD>cpa dsRNA* are also shown in Fig. 6E and G. Statistical analysis as described in Fig. 6.

(M) Comparison of SPY-labeled free cable growth in Flr KD using two independent transgenes. Control and *MTD>flr dsRNA (1)* data are also shown in Fig. 5G and H. Statistical analysis as described in Fig. 5.

**Figure S2: Patronin and Shot KD produces free cables.** (A,B) Representative images of fixed s10B.4 nurse cells triple-labeled for actin cables (grey; phallodin), DNA (DAPI; grey or cyan) and the Ace-MT network (magenta; anti-Ace-α-tubulin) from (A) *MTD>patronin dsRNA* and (B) *MTD>shot^ROD^ dsRNA.* These are shown at lower magnification (scale bar = 10 µm) and higher magnification (scale bar = 5 µm). Images were taken of the same nurse cells for the lower magnification views and the higher magnification views of actin and cytoplasmic Ace-MTs. Independent, stage-matched nurse cells were used for the images of the cortical Ace-MT network. The cortical Ace-MT network is shown *en face* and all other images are cross-sections. Yellow arrowheads indicate free cables. (C) Quantification of the proportion of cortical versus free cables during s10B in genotypes as shown. Chi-square test for goodness fit, (2) is the expected and compared to (3-6), (7) is the expected and compared to (8). Conditions sharing letters are not significantly different from one another, conditions with different letters are significantly different p<0.0001, n=20-45 egg chambers per genotype with all substages represented.

**Figure S3: Live imaging of cables from representative genotypes/conditions.** SiR- or SPY-labeled actin cables in genotypes/conditions as shown. In all cases, images were taken every 15 min. Scale bar = 3 µm. The cortex is indicated with a white dashed line in images where it isn’t visible in that focal plane.

**Figure S4: Representative images from FRAP of GFP-actin cortex and distal regions of GFP-actin cables** (A-A5) Representative images of live nurse cells expressing GFP-Act5C with photobleaching at the cortex and the proximal end of the cable. (A) Pre-bleaching. The region to be bleached is marked in yellow. (A0) shows post-bleached cortex and cable. (A1-A5) shows time points over 15 minutes after photobleaching. Both the actin cortex and the proximal cable show recovery. See Fig. 7K for quantification of fluorescence recovery. Scale bar = 3 µm

(B-B3) Representative images of live nurse cells expressing GFP-Act5C with photobleaching in a distal region of the cable. (B) Pre-bleaching. The region to be bleached is marked in yellow. (B0) shows a post-bleached cable. (B1-B3) shows time points over 15 minutes after photobleaching. The distal bleached region of the cable did not show significant recovery and moved away from the cortex as the cable elongated. See Fig. 7K for quantification of fluorescence recovery. Scale bar = 3 µm

**Figure S5:**

(A) Immunofluorescent images (max projection of four 0.4 µm z-slices) of WT nurse cells in cross section at 10B.4 triple-labeled for Actin (grey; phalloidin), Capping Protein (Cpa, cyan) and Ena (green) (scale bar = 5 µm).

(B) Quantification of cortical Ace-MT density in WT and *MDD>cpa dsRNA* nurse cells in stages as shown. One-way ANOVA with Tukey’s multiple comparison test. ****p<0.0001, n=5-12 per substage.

(C,D) Images (max projection of fourteen 0.4 µm z-slices) from fixed s9 nurse cells in cross section double labeled for actin cables (grey; phallodin) and the Ace-MT network (magenta; anti-Ace-α-tubulin) from control (C, *MDD>*) and *MDD>cpa dsRNA* (D) genotypes. (scale bar = 5 µm)

(E-G) Images of s10B actin cables in live nurse cells labeled with SPY-actin. Actin cables were imaged for 90 minutes; representative time points at t=0, t=30, and t=60 minutes are shown. The cortex is indicated with white dashed lines or white arrowheads, and a representative cable is indicated with yellow arrowheads. (scale bar = 3 µm)

(H) Quantification of cable length over time for genotypes and conditions shown. Initial cable lengths were normalized to 0 µm at time = 0:00 for comparison. In WT+DMSO treated control cells, elongating cortical cables were measured. In KDs and drug-treated conditions, cables detach at time zero and free cable lengths were measured. Genotypes sharing letters are not statistically different from one another. p<0.0001. Simple linear regression with ANOVA. n=20-25 cables per genotype/condition.

## References

Alimov N, Hoeprich GJ, Padrick SB, Goode BL. (2023) Cyclase-associated protein interacts with actin filament barbed ends to promote depolymerization and formin displacement. J Biol Chem. Dec;299(12):105367. doi: 10.1016/j.jbc.2023.105367. Epub 2023 Oct 19. PMID: 37863260; PMCID: PMC10692737.

Imran Alsous J, Romeo N, Jackson JA, Mason FM, Dunkel J, Martin AC. (2021) Dynamics of hydraulic and contractile wave-mediated fluid transport during *Drosophila* oogenesis. Proc Natl Acad Sci U S A. Mar 9;118(10):e2019749118. doi: 10.1073/pnas.2019749118. PMID: 33658367; PMCID: PMC7958293.

Arber, S., Barbayannis, F. A., Hanser, H., Schneider, C., Stanyon, C. A., Bernard, O., & Caroni, P. (1998). Regulation of actin dynamics through phosphorylation of cofilin by LIM-kinase. Nature, 393(6687), 805–809. 10.1038/31729

Arya A, Choubey S, Shekhar S. (2024) Actin Filament Barbed-End Depolymerization by Combined Action of Profilin, Cofilin, and Twinfilin. PRX Life. Jul-Sep;2(3):033002. doi: 10.1103/prxlife.2.033002. Epub 2024 Jul 16. PMID: 40777877; PMCID: PMC12330814.

Avenarius MR, Krey JF, Dumont RA, Morgan CP, Benson CB, Vijayakumar S, Cunningham CL, Scheffer DI, Corey DP, Müller U, Jones SM, Barr-Gillespie PG. (2017) Heterodimeric capping protein is required for stereocilia length and width regulation. J Cell Biol. Nov 6;216(11):3861–3881. doi: 10.1083/jcb.201704171. Epub 2017 Sep 12. PMID: 28899994; PMCID: PMC5674897.

Bieling, P., Kandels-Lewis, S., Telley, I. A., van Dijk, J., Janke, C., & Surrey, T. (2008). CLIP- 170 tracks growing microtubule ends by dynamically recognizing composite EB1/tubulinbinding sites. Journal of Cell Biology, 183(7), 1223–1233. 10.1083/jcb.200809190

Bilancia, C. G., Winkelman, J. D., Tsygankov, D., Nowotarski, S. H., Sees, J. A., Comber, K., Evans, I., Lakhani, V., Wood, W., Elston, T. C. et al. (2014). Enabled negatively regulates diaphanous-driven actin dynamics in vitro and in vivo. Dev. Cell 28, 394–408. doi:10.1016/j.devcel.2014.01.015

Blanchoin, L., Boujemaa-Paterski, R., Sykes, C., & Plastino, J. (2014). Actin dynamics, architecture, and mechanics in cell motility. Physiological Reviews, 94(1), 235–263.

Bombardier JP, Eskin JA, Jaiswal R, Corrêa IR Jr, Xu MQ, Goode BL, Gelles J. (2015) Single-molecule visualization of a formin-capping protein ‘decision complex’ at the actin filament barbed end. Nat Commun. Nov 13;6:8707. doi: 10.1038/ncomms9707. PMID: 26566078; PMCID: PMC4660045.

Breitsprecher D, Koestler SA, Chizhov I, Nemethova M, Mueller J, Goode BL, Small JV, Rottner K, Faix J. (2011) Cofilin cooperates with fascin to disassemble filopodial actin filaments. J Cell Sci. Oct 1;124(Pt 19):3305–18. doi: 10.1242/jcs.086934. PMID: 21940796; PMCID: PMC4074248.

Bugyi B, Carlier MF. (2010) Control of actin filament treadmilling in cell motility. Annu Rev Biophys.;39:449–70. doi: 10.1146/annurev-biophys-051309-103849. PMID: 20192778.

Cant, K., Knowles, B. A., Mooseker, M. S., & Cooley, L. (1994). Drosophila singed, a fascin homolog, is required for actin bundle formation during oogenesis and bristle extension. Journal of Cell Biology, 125(2), 369–380.

Caldwell JE, Heiss SG, Mermall V, Cooper JA. (1989) Effects of CapZ, an actin capping protein of muscle, on the polymerization of actin. Biochemistry. Oct 17;28(21):8506-14. doi: 10.1021/bi00447a036. PMID: 2557904.

Chikireddy, J., Lengagne, L., Le Borgne, R., Durieu, C., Wioland, H., Romet-Lemonne, G., & J.gou, A. (2024). Fascin-induced bundling protects actin filaments from disassembly by cofilin. Journal of Cell Biology, 223(6), e202312106.

Chou WC, Lakonishok M, Lu W, Gelfand VI, McCartney BM. (2026) Developmentally regulated actin-microtubule cross talk in Drosophila oogenesis. J Cell Biol. Mar 2;225(3):e202411007. doi: 10.1083/jcb.202411007. Epub 2026 Jan 9. PMID: 41511043.

Chugh P, Clark AG, Smith MB, Cassani DAD, Dierkes K, Ragab A, Roux PP, Charras G, Salbreux G, Paluch EK. (2017) Actin cortex architecture regulates cell surface tension. Nat Cell Biol. Jun;19(6):689–697. doi: 10.1038/ncb3525. Epub 2017 May 22. PMID: 28530659; PMCID: PMC5536221.

Coue, M., Brenner, S. L., Spector, I., & Korn, E. D. (1987). Inhibition of actin polymerization by latrunculin A. FEBS Letters, 213(2), 316–318. 10.1016/0014-5793(87)81513-2

Courtemanche N, Henty-Ridilla JL. (2024) Actin filament dynamics at barbed ends: New structures, new insights. Curr Opin Cell Biol. Oct;90:102419. doi: 10.1016/j.ceb.2024.102419. Epub 2024 Aug 22. PMID: 39178734; PMCID: PMC11492572.

Delalle I, Pfleger CM, Buff E, Lueras P, Hariharan IK. (2005) Mutations in the Drosophila orthologs of the F-actin capping protein alpha- and beta-subunits cause actin accumulation and subsequent retinal degeneration. Genetics. Dec;171(4):1757–65. doi: 10.1534/genetics.105.049213. Epub 2005 Sep 2. PMID: 16143599; PMCID: PMC1456101.

Drummond MC, Barzik M, Bird JE, Zhang DS, Lechene CP, Corey DP, Cunningham LL, Friedman TB. (2015) Live-cell imaging of actin dynamics reveals mechanisms of stereocilia length regulation in the inner ear. Nat Commun. Apr 21;6:6873. doi: 10.1038/ncomms7873. PMID: 25898120; PMCID: PMC4411292.

Eshun-Wilson, L., Zhang, R., Portran, D., Nachury, M. V., Toso, D. B., L.hr, T., Vendruscolo, M., Bonomi, M., Fraser, J. S., & Nogales, E. (2019). Effects of α-tubulin acetylation on microtubule structure and stability. Proceedings of the National Academy of Sciences, 116(21), 10366–10371. 10.1073/pnas.1900441116

Folker, E. S., Baker, B. M., & Goodson, H. V. (2005). Interactions between CLIP-170, Tubulin, and Microtubules: Implications for the Mechanism of CLIP-170 Plus-End Tracking Behavior. Molecular Biology of the Cell, 16(11), 5373–5384.

Fritzsche M, Lewalle A, Duke T, Kruse K, Charras G. (2013) Analysis of turnover dynamics of the submembranous actin cortex. Mol Biol Cell. Mar;24(6):757–67. doi: 10.1091/mbc.E12-06-0485. Epub 2013 Jan 23. PMID: 23345594; PMCID: PMC3596247.

Gates J, Mahaffey JP, Rogers SL, Emerson M, Rogers EM, Sottile SL, Van Vactor D, Gertler FB, Peifer M. (2007) Enabled plays key roles in embryonic epithelial morphogenesis in Drosophila. Development. Jun;134(11):2027–39. doi: 10.1242/dev.02849. PMID: 17507404.

Gates, J., Nowotarski, S. H., Yin, H., Mahaffey, J. P., Bridges, T., Herrera, C., Homem, C. C. F., Janody, F., Montell, D. J., & Peifer, M. (2009). Enabled and Capping protein play important roles in shaping cell behavior during Drosophila oogenesis. Developmental Biology, 333(1), 90–107.

Goode, B. L., Drubin, D. G., & Lappalainen, P. (1998). Regulation of the cortical actin cytoskeleton in budding yeast by twinfilin, a ubiquitous actin monomer-sequestering protein. The Journal of Cell Biology, 142(3), 723–733. 10.1083/jcb.142.3.723

Goode, B. L., Eskin, J., & Shekhar, S. (2023). Mechanisms of actin disassembly and turnover. Journal of Cell Biology, 222(12), e202309021.

Guild, G. M., Connelly, P. S., Shaw, M. K., & Tilney, L. G. (1997). Actin Filament Cables in Drosophila Nurse Cells Are Composed of Modules That Slide Passively Past One Another during Dumping. The Journal of Cell Biology, 138(4), 783–797.

Hakala M, Wioland H, Tolonen M, Kotila T, Jegou A, Romet-Lemonne G, Lappalainen P. (2021) Twinfilin uncaps filament barbed ends to promote turnover of lamellipodial actin networks. Nat Cell Biol. Feb;23(2):147–159. doi: 10.1038/s41556-020-00629-y. Epub 2021 Feb 8. Erratum in: Nat Cell Biol. 2021 Apr;23(4):437-438. doi: 10.1038/s41556-021-00651-8. PMID: 33558729.

Harker, A. J., Katkar, H. H., Bidone, T. C., Aydin, F., Voth, G. A., Applewhite, D. A., & Kovar, D. R. (2019). Ena/VASP processive elongation is modulated by avidity on actin filaments bundled by the filopodia cross-linker fascin. Molecular Biology of the Cell, 30(7), 851– 862. 10.1091/mbc.E18-08-0500

Henty-Ridilla, J. L., Rankova, A., Eskin, J. A., Kenny, K., & Goode, B. L. (2016). Accelerated actin filament polymerization from microtubule plus ends. Science.

Hopmann R, Cooper JA, Miller KG. (1996) Actin organization, bristle morphology, and viability are affected by actin capping protein mutations in Drosophila. J Cell Biol. Jun;133(6):1293–305. doi: 10.1083/jcb.133.6.1293. PMID: 8682865; PMCID: PMC2120907.

Huelsmann, S., Yl.nne, J., & Brown, N. H. (2013). Filopodia-like Actin Cables Position Nuclei in Association with Perinuclear Actin in Drosophila Nurse Cells. Developmental Cell, 26(6), 604–615.

Janody F, Treisman JE. (2006) Actin capping protein alpha maintains vestigial-expressing cells within the Drosophila wing disc epithelium. Development. Sep;133(17):3349–57. doi: 10.1242/dev.02511. Epub 2006 Aug 3. PMID: 16887822; PMCID: PMC1544359.

Jayo A, Parsons M, Adams JC. A novel Rho-dependent pathway that drives interaction of fascin-1 with p-Lin-11/Isl-1/Mec-3 kinase (LIMK) 1/2 to promote fascin-1/actin binding and filopodia stability. BMC Biol. 2012 Aug 10;10:72. doi: 10.1186/1741-7007-10-72. PMID: 22883572; PMCID: PMC3488970.

Johnston AB, Collins A, Goode BL. (2015) High-speed depolymerization at actin filament ends jointly catalysed by Twinfilin and Srv2/CAP. Nat Cell Biol. Nov;17(11):1504–11. doi: 10.1038/ncb3252. Epub 2015 Oct 12. PMID: 26458246; PMCID: PMC4808055.

Kachar, B. (2019). Moving Encounters: Actin Treadmilling in the Brush Border. Developmental Cell, 50(5), 529–530.

Kelkar M, Bohec P, Charras G. (2020) Mechanics of the cellular actin cortex: From signalling to shape change. Curr Opin Cell Biol. Oct;66:69–78. doi: 10.1016/j.ceb.2020.05.008. Epub 2020 Jun 21. PMID: 32580115.

Kellogg, E. H., Hejab, N. M. A., Howes, S., Northcote, P., Miller, J. H., D.az, J. F., Downing, K. H., & Nogales, E. (2017). Insights into the Distinct Mechanisms of Action of Taxane and Non-Taxane Microtubule Stabilizers from Cryo-EM Structures. Journal of Molecular Biology, 429(5), 633–646. 10.1016/j.jmb.2017.01.001

Kemp JP Jr, Brieher WM. (2018) The actin filament bundling protein α-actinin-4 actually suppresses actin stress fibers by permitting actin turnover. J Biol Chem. Sep 14;293(37):14520-14533. doi: 10.1074/jbc.RA118.004345. Epub 2018 Jul 26. PMID: 30049798; PMCID: PMC6139541.

Komarova Y, De Groot CO, Grigoriev I, Gouveia SM, Munteanu EL, Schober JM, Honnappa S, Buey RM, Hoogenraad CC, Dogterom M, Borisy GG, Steinmetz MO, Akhmanova A. (2009) Mammalian end binding proteins control persistent microtubule growth. J Cell Biol. Mar 9;184(5):691–706. doi: 10.1083/jcb.200807179. Epub 2009 Mar 2. PMID: 19255245; PMCID: PMC2686402.

Lappalainen P. (2016) Actin-binding proteins: the long road to understanding the dynamic landscape of cellular actin networks. Mol Biol Cell. Aug 15;27(16):2519-22. doi: 10.1091/mbc.E15-10-0728. PMID: 27528696; PMCID: PMC4985253.

Lewkowicz, E., Herit, F., Le Clainche, C., Bourdoncle, P., Perez, F., & Niedergang, F. (2008). The microtubule-binding protein CLIP-170 coordinates mDia1 and actin reorganization during CR3-mediated phagocytosis. Journal of Cell Biology, 183(7), 1287–1298. 10.1083/jcb.200807023

Logan, G., & McCartney, B. (2020). Comparative analysis of taxol-derived fluorescent probes to assess microtubule networks in a complex live three-dimensional tissue. Cytoskeleton, 77(5–6), 229–237. 10.1002/cm.21599

Logan, G., Chou, W.-C., & McCartney, B. M. (2022). A Diaphanous and Enabled-dependent asymmetric actin cable array repositions nuclei during Drosophila oogenesis. Development (Cambridge, England), 149(13), dev197442. 10.1242/dev.197442

Maurer, S. P., Fourniol, F. J., Bohner, G., Moores, C. A., & Surrey, T. (2012). EBs recognize a nucleotide-dependent structural cap at growing microtubule ends. Cell, 149(2), 371–382. 10.1016/j.cell.2012.02.049

Mahajan-Miklos, S., & Cooley, L. (1994). Intercellular cytoplasm transport during Drosophila oogenesis. Developmental Biology, 165(2), 336–351.

Matova, N., Mahajan-Miklos, S., Mooseker, M. S., & Cooley, L. (1999). Drosophila Quail, a villin related protein, bundles actin filaments in apoptotic nurse cells. Development, 126(24), 5645–5657. 10.1242/dev.126.24.5645

Meenderink, L. M., Gaeta, I. M., Postema, M. M., Cencer, C. S., Chinowsky, C. R., Krystofiak, E. S., Millis, B. A., & Tyska, M. J. (2019). Actin dynamics drive microvillar motility and clustering during brush border assembly. Developmental Cell, 50(5), 545–556.e4.

Mejillano MR, Kojima S, Applewhite DA, Gertler FB, Svitkina TM, Borisy GG. (2004) Lamellipodial versus filopodial mode of the actin nanomachinery: pivotal role of the filament barbed end. Cell. Aug 6;118(3):363-73. doi: 10.1016/j.cell.2004.07.019. PMID: 15294161.

Michelot, A., & Drubin, D. G. (2011). Building distinct actin filament networks in a common cytoplasm. Current Biology: CB, 21(14), R560–569.

Mitani T, Takeda S, Oda T, Narita A, Maéda Y, Honda H, Fujiwara I. (2025) Microscopic and structural observations of actin filament capping and severing by cytochalasin D. Proc Natl Acad Sci U S A. 2025 Jul 22;122(29):e2502164122. doi: 10.1073/pnas.2502164122. Epub 2025 Jul 14. Erratum in: Proc Natl Acad Sci U S A. 2025 Sep 23;122(38):e2523366122. doi: 10.1073/pnas.2523366122. PMID: 40658853; PMCID: PMC12304888.

Mogilner, A., & Rubinstein, B. (2005). The Physics of Filopodial Protrusion. Biophysical Journal, 89(2), 782–795.

Narayanan, P., Chatterton, P., Ikeda, A., Ikeda, S., Corey, D. P., Ervasti, J. M., & Perrin, B. J. (2015). Length regulation of mechanosensitive stereocilia depends on very slow actin dynamics and filament-severing proteins. Nature Communications, 6(1), 6855.

Perez, F., Diamantopoulos, G. S., Stalder, R., & Kreis, T. E. (1999). CLIP-170 Highlights Growing Microtubule Ends In Vivo. Cell, 96(4), 517–527. 10.1016/S0092-8674(00)80656-X

Piehl, M., & Cassimeris, L. (2003). Organization and Dynamics of Growing Microtubule Plus Ends during Early Mitosis. Molecular Biology of the Cell, 14(3), 916–925. 10.1091/mbc.e02-09-0607

Pollard, T. D. (2016). Actin and Actin-Binding Proteins. Cold Spring Harbor Perspectives in Biology, 8(8), a018226. 10.1101/cshperspect.a018226

Rajan S, Kudryashov DS, Reisler E. (2023) Actin Bundles Dynamics and Architecture. Biomolecules. Feb 28;13(3):450. doi: 10.3390/biom13030450. PMID: 36979385; PMCID: PMC10046292.

Reddy V, Arya A, Shekhar S. (2025) Twinfilin is a nonprocessive depolymerase which synergizes with formin to dramatically accelerate actin filament uncapping by 300-fold. Proc Natl Acad Sci U S A. May 6;122(18):e2501078122. doi: 10.1073/pnas.2501078122. Epub 2025 Apr 28. PMID: 40294253; PMCID: PMC12067289.

Rizvi SA, Neidt EM, Cui J, Feiger Z, Skau CT, Gardel ML, Kozmin SA, Kovar DR. (2009) Identification and characterization of a small molecule inhibitor of formin-mediated actin assembly. Chem Biol. Nov 25;16(11):1158-68. doi: 10.1016/j.chembiol.2009.10.006. PMID: 19942139; PMCID: PMC2784894.

Rodal, A. A., Tetreault, J. W., Lappalainen, P., Drubin, D. G., & Amberg, D. C. (1999). Aip1p interacts with cofilin to disassemble actin filaments. The Journal of Cell Biology, 145(6), 1251–1264. 10.1083/jcb.145.6.1251

Romet-Lemonne G, Jégou A. (2021) The dynamic instability of actin filament barbed ends. J Cell Biol. Apr 5;220(4):e202102020. doi: 10.1083/jcb.202102020. PMID: 33755041; PMCID: PMC8103424.

Rzadzinska AK, Nevalainen EM, Prosser HM, Lappalainen P, Steel KP. (2009) MyosinVIIa interacts with Twinfilin-2 at the tips of mechanosensory stereocilia in the inner ear. PLoS One. Sep 23;4(9):e7097. doi: 10.1371/journal.pone.0007097. PMID: 19774077; PMCID: PMC2743196.

Shekhar S, Hoeprich GJ, Gelles J, Goode BL. (2021) Twinfilin bypasses assembly conditions and actin filament aging to drive barbed end depolymerization. J Cell Biol. Jan 4;220(1):e202006022. doi: 10.1083/jcb.202006022. PMID: 33226418; PMCID: PMC7686915.

Sherer, L. A., & Courtemanche, N. (2022). Cooperative bundling by fascin generates actin structures with architectures that depend on filament length. Frontiers in Cell and Developmental Biology, 10, 974047. 10.3389/fcell.2022.974047

Tilney, L. G., Derosier, D. J., & Mulroy, M. J. (1980). The organization of actin filaments in the stereocilia of cochlear hair cells. Journal of Cell Biology, 86(1), 244–259.

Tilney LG, Tilney MS, Guild GM. (1995) F actin bundles in Drosophila bristles. I. Two filament cross-links are involved in bundling. J Cell Biol. Aug;130(3):629–38. doi: 10.1083/jcb.130.3.629. PMID: 7622563; PMCID: PMC2120540.

Tilney LG, Connelly PS, Ruggiero L, Vranich KA, Guild GM. (2003) Actin filament turnover regulated by cross-linking accounts for the size, shape, location, and number of actin bundles in Drosophila bristles. Mol Biol Cell. Oct;14(10):3953–66. doi: 10.1091/mbc.e03-03-0158. Epub 2003 Jul 25. PMID: 14517310; PMCID: PMC206991.

Tyska MJ, Mooseker MS. (2002) MYO1A (brush border myosin I) dynamics in the brush border of LLC-PK1-CL4 cells. Biophys J. Apr;82(4):1869–83. doi: 10.1016/S0006-3495(02)75537-9. PMID: 11916846; PMCID: PMC1301984.

Ulrichs, H., Gaska, I., & Shekhar, S. (2023). Multicomponent regulation of actin barbed end assembly by twinfilin, formin and capping protein. Nature Communications, 14(1),

Wahlstrom, G., Vartiainen, M., Yamamoto, L., Mattila, P. K., Lappalainen, P., & Heino, T. I. (2001). Twinfilin is required for actin-dependent developmental processes in Drosophila. Journal of Cell Biology, 155(5), 787–796. 10.1083/jcb.200108022

Wear MA, Yamashita A, Kim K, Maéda Y, Cooper JA. (2003) How capping protein binds the barbed end of the actin filament. Curr Biol. Sep 2;13(17):1531-7. doi: 10.1016/s0960-9822(03)00559-1. PMID: 12956956.

Winkelman, J. D., Bilancia, C. G., Peifer, M., & Kovar, D. R. (2014). Ena/VASP Enabled is a highly processive actin polymerase tailored to self-assemble parallel-bundled F-actin networks with Fascin. Proceedings of the National Academy of Sciences of the United States of America, 111(11), 4121–4126. 10.1073/pnas.1322093111

Wioland H, Guichard B, Senju Y, Myram S, Lappalainen P, Jégou A, Romet-Lemonne G. (2017) ADF/Cofilin Accelerates Actin Dynamics by Severing Filaments and Promoting Their Depolymerization at Both Ends. Curr Biol. Jul 10;27(13):1956–;.e7. doi: 10.1016/j.cub.2017.05.048. Epub 2017 Jun 15. PMID: 28625781; PMCID: PMC5505867.

Wioland H, Jegou A, Romet-Lemonne G. (2019) Quantitative Variations with pH of Actin Depolymerizing Factor/Cofilin’s Multiple Actions on Actin Filaments. Biochemistry. Jan 8;58(1):40–47. doi: 10.1021/acs.biochem.8b01001. Epub 2018 Dec 18. PMID: 30499293; PMCID: PMC6358128.

Wirshing ACE, Rodriguez SG, Goode BL. (2023) Evolutionary tuning of barbed end competition allows simultaneous construction of architecturally distinct actin structures. J Cell Biol. Apr 3;222(4):e202209105. doi: 10.1083/jcb.202209105. Epub 2023 Feb 2. PMID: 36729023; PMCID: PMC9929936.

Yang, N., Higuchi, O., Ohashi, K., Nagata, K., Wada, A., Kangawa, K., Nishida, E., & Mizuno, K. (1998). Cofilin phosphorylation by LIM-kinase 1 and its role in Rac-mediated actin reorganization. Nature, 393(6687), Article 6687. 10.1038/31735

