## Supplemental Tables and Figures for "Microtubules coordinate with actin assembly and disassembly to elongate cortical actin cables"

**Table 1:** Summary of protein depletion/knockdown efficiency and validation of phenotypes with additional genotypes (also see Fig. S1).

| Protein | Verification & degree of depletion | Phenotype verification |
|---|---|---|
| CLIP-190 | RT-PCR of whole ovaries from *MTD>CLIP-190 dsRNA* (1) and (2) showed an approx. 50% reduction in mRNA (Fig. S1A,B). | We examined cytoplasmic MT density, cortical cable density, cable length, and cortical vs free cables for both transgenes (FIg. S1G-J). |
| Eb1 | RT-PCR of whole ovaries from *MTD>Eb1 dsRNA* showed an approx. 50% reduction in mRNA (Fig. S1A,C). | We examined cytoplasmic MT density, cortical cable density, cable length, and cortical vs free cables for the Eb1 dsRNA and an Eb1 dominant negative (DN) (FIg. S1G-J). |
| $\gamma$-tubulin | Previously shown: $\gamma$-tubulin antibody, quantified number of $\gamma$-tubulin puncta (Chou et al., 2026) | We examined cytoplasmic MT density, cortical cable density, and cable length for both transgenes (FIg. S1G-J). |
| Patronin | Previously shown: Patronin antibody, quantified cortical to cytoplasmic ratio of fluorescence intensity for *UASp-Patronin dsRNA* (Chou et al., 2026) | In addition to *UASp-patonin dsRNA,* we expressed a *UASt-patronin dsRNA* in the germline by simultaneously knocking down *Rhino* expression and compared this to *Rhino* knock down alone for the frequency of free cables (Fig. S2C). *Rhino* knock down enables UASt-based transgenes to be expressed during oogenesis (Deluca & Spradling et al., 2018). |
| Shot | Previously shown: Shot antibody, cortical to cytoplasmic ratio of fluorescence intensity (Chou et al., 2026) | We used a dsRNA that targets all 22 Shot isoforms through the Rod domain (*shot$^{Rod}$*) and a dsRNA that specifically targets the 14 ABD-containing isoforms (*shot$^{ABD}$*) (Lu et al., 2021). Both have comparable increases in the frequency of free cables (Fig. S2C). |
| Dia | Previously shown: using Dia antibody and quantifying the cortical to cytoplasmic ratio of fluorescence intensity demonstrating an approx. 50-60% reduction in Dia protein with both dsRNA transgenes (Logan et al., 2022). | We previously reported a comparable defect in actin cable density and no change to cytoplasmic MT density for two independent *dia dsRNA* transgenes (Chou et al., 2026). |
| Ena | Previously shown: using Ena antibody and quantifying the reduction in Ena protein by Western blot. This showed a >50% reduction in Ena (Logan et al., 2022). | We previously reported a comparable defect in actin cable density and no change to cytoplasmic MT density for two independent *ena dsRNA* transgenes (Chou et al., 2026). |
| Quail | Previously shown: Transheterozygous *quail* mutant using two EMS-induced alleles (*qua$^1$/qua$^{6-396}$*). *qua$^1$* exhibits a significantly reduced mRNA level (Mahajan-Miklos and Cooley, 1994). | We previously reported a comparable defect in actin cable density and cytoplasmic MT density for *qua$^1$/qua$^{6-396}$* and for a *qua dsRNA* transgene (Chou et al., 2026). |
| Singed | Previously shown: The homozygous loss-of-function allele (*sn$^{x2}$/sn$^{x2}$*) produced no detectable protein (Cant et al., 1994). | We previously reported a comparable defect in actin cable density and cytoplasmic MT density for *sn$^{x2}$/sn$^{x2}$* and for a *sn dsRNA* transgene (Chou et al., 2026). |
| Fimbrin | Previously shown: *Fimbrin* dsRNA expression significantly reduced Fimbrin-EGFP shown by the cortical to cytoplasmic ratio of fluorescence intensity (Chou et al., 2026). | We previously reported a comparable defect in actin cable density and cytoplasmic MT density two independent *Fim dsRNA* transgenes (Chou et al., 2026). |
| LIMK | RT-PCR of whole ovaries from *MTD>LIMK dsRNA* (1) showed an approx. 25% reduction in mRNA (Fig. S1A,D). | We examined cortical cable density and cytoplasmic MT density for both *LIMK dsRNA* transgenes (Fig. S1K,L). |
| Flare | RT-PCR of whole ovaries from *MTD>flr dsRNA* (1) showed an approx. 50% reduction in mRNA (Fig. S1A,E). | We observed free cable elongation in cells from both *flr dsRNA* transgenes (Fig. S1M). |
| Capping Protein | Using CP antibody, we assessed the cortical to cytoplasmic ratio of fluorescence intensity for *MDD>cpa dsRNA* (FIg. S1F). | We examined cortical cable density and cytoplasmic MT density for both *cpa* and *cpb dsRNA* transgenes (Fig. S1K,L). |

| TABLE 2 | Microtubule +Tip proteins | | | Microtubule nucleation/tethering | | | Microtubule drugs | | Actin assembly factors | | Actin filament bundlers | | | Actin filament turnover regulation | | | | | | |
|---|---|---|---|---|---|---|---|---|---|---|---|---|---|---|---|---|---|---|---|---|
| | **CLIP KD** | **CLIP KD + CytoD** | **EB1 KD** | **γ-tubulin KD** | **Patronin KD** | **Shot KD** | **WT + nocodazole** | **WT + taxol** | **Dia KD** | **Ena KD** | **Quail mutant** | **Singed mutant** | **Fimbrin KD** | **WT + LatA** | **LIMK KD** | **Flare KD** | **Flr KD + SMIFH2** | **WT + CytoD** | **Cpa KD** | **Cpa KD + CytoD** |
| **manipulation** | loss of MT regulation | loss of MT regulation + barbed-end capping | loss of MT regulation | reduced MT nucleation | reduced MT minus end binding | reduced cortical tethering of MTs | depletion of MT network | stabilization of MTs | reduced barbed end assembly | reduced barbed end assembly | reduced filament bundling | reduced filament bundling | reduced filament bundling | monomer binding-reduced barbed-end assembly | increased Cofilin activity | reduced Cofilin activity | reduced Cofilin-mediated disassembly and reduced formin-mediated assembly | barbed end capping- block barbed end assembly and disassembly | reduced barbed end capping | recapping the barbed end |
| **Cytoplasmic MT density** | reduced | nd | reduced | reduced | reduced* | reduced* | reduced* | nd | wild type* | wild type* | reduced* | reduced* | reduced* | reduced | reduced | wildtype | nd | nd | reduced | nd |
| **Cortical MT density** | reduced | nd | reduced | reduced | reduced* | reduced* | reduced* | nd | reduced* | reduced* | reduced* | reduced* | reduced* | reduced | reduced | wildtype | nd | nd | reduced | nd |
| **Cortical actin cable density** | reduced | nd | reduced | reduced | reduced* | reduced* | nd | nd | reduced* | reduced* | reduced* | reduced* | reduced* | nd | reduced | reduced | nd | nd | reduced | nd |
| **Cortical actin cable length** | reduced | nd | reduced | reduced | reduced* | reduced* | nd | nd | reduced* | reduced* | reduced* | reduced* | reduced* | nd | reduced | wild type | nd | nd | reduced | nd |
| **Free actin cables** | increased | yes | increased | increased | increased | increased | yes | yes | wild type | wild type | reduced | reduced | reduced | increased | increased | increased | increased | yes | increased | increased |
| **Free cable behavior** | shrink | stable length | shrink | stable length | nd | nd | stable length | stable length | na | na | na | na | na | shrink | shrink | grow | stable length | stable length | shrink | stable length |
| **Cortical actin cable-MT coalignment** | reduced | nd | reduced | increased | nd | nd | nd | nd | wild type* | wild type* | increased* | increased* | increased* | nd | increased | wild type | nd | nd | increased | nd |
| **Free cable-MT coalignment** | wild type | nd | nd | increased | nd | nd | nd | nd | na | na | na | na | na | nd | increased | increased | nd | nd | increased | nd |

### Figure S1

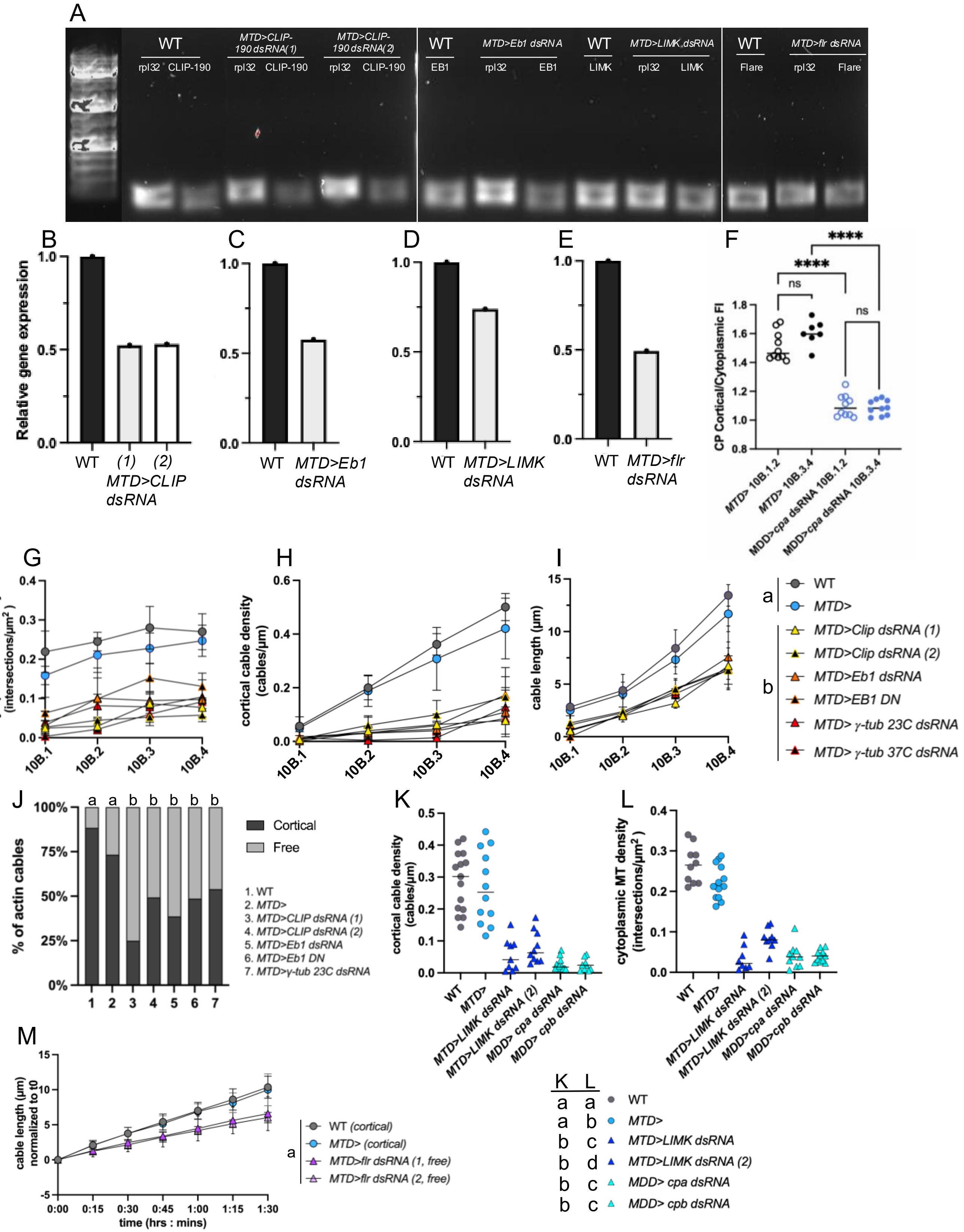

A
WT
MTD>CLIP-190 dsRNA(1)
MTD>CLIP-190 dsRNA(2)
rpl32 CLIP-190
WT
MTD>Eb1 dsRNA
EB1
rpl32 EB1
WT
MTD>LIMK dsRNA
LIMK
rpl32 LIMK
WT
MTD>flr dsRNA
Flare
rpl32 Flare
B
C
D
E
F
Relative gene expression
WT (1) (2) MTD>CLIP dsRNA
WT MTD>Eb1 dsRNA
WT MTD>LIMK dsRNA
WT MTD>flr dsRNA
CP Cortical/Cytoplasmic FI
MTD> 10B.1.2
MTD> 10B.3.4
MDD>cpa dsRNA 10B.1.2
MDD>cpa dsRNA 10B.3.4
ns
****
G
H
I
cytoplasmic MT density (intersections/μm²)
cortical cable density (cables/μm)
cable length (μm)
10B.1 10B.2 10B.3 10B.4
a
WT
MTD>
b
MTD>Clip dsRNA (1)
MTD>Clip dsRNA (2)
MTD>Eb1 dsRNA
MTD>EB1 DN
MTD> γ-tub 23C dsRNA
MTD> γ-tub 37C dsRNA
J
% of actin cables
a a b b b b b
Cortical
Free
1. WT
2. MTD>
3. MTD>CLIP dsRNA (1)
4. MTD>CLIP dsRNA (2)
5. MTD>Eb1 dsRNA
6. MTD>Eb1 DN
7. MTD>γ-tub 23C dsRNA
K
L
cortical cable density (cables/μm)
cytoplasmic MT density (intersections/μm²)
WT
MTD>
MTD>LIMK dsRNA
MTD>LIMK dsRNA (2)
MDD> cpa dsRNA
MDD> cpb dsRNA
M
cable length (μm) normalized to t0
time (hrs : mins)
a
WT (cortical)
MTD> (cortical)
MTD>flr dsRNA (1, free)
MTD>flr dsRNA (2, free)
K L
a a WT
a b MTD>
b c MTD>LIMK dsRNA
b d MTD>LIMK dsRNA (2)
b c MDD> cpa dsRNA
b c MDD> cpb dsRNA

### Figure S2

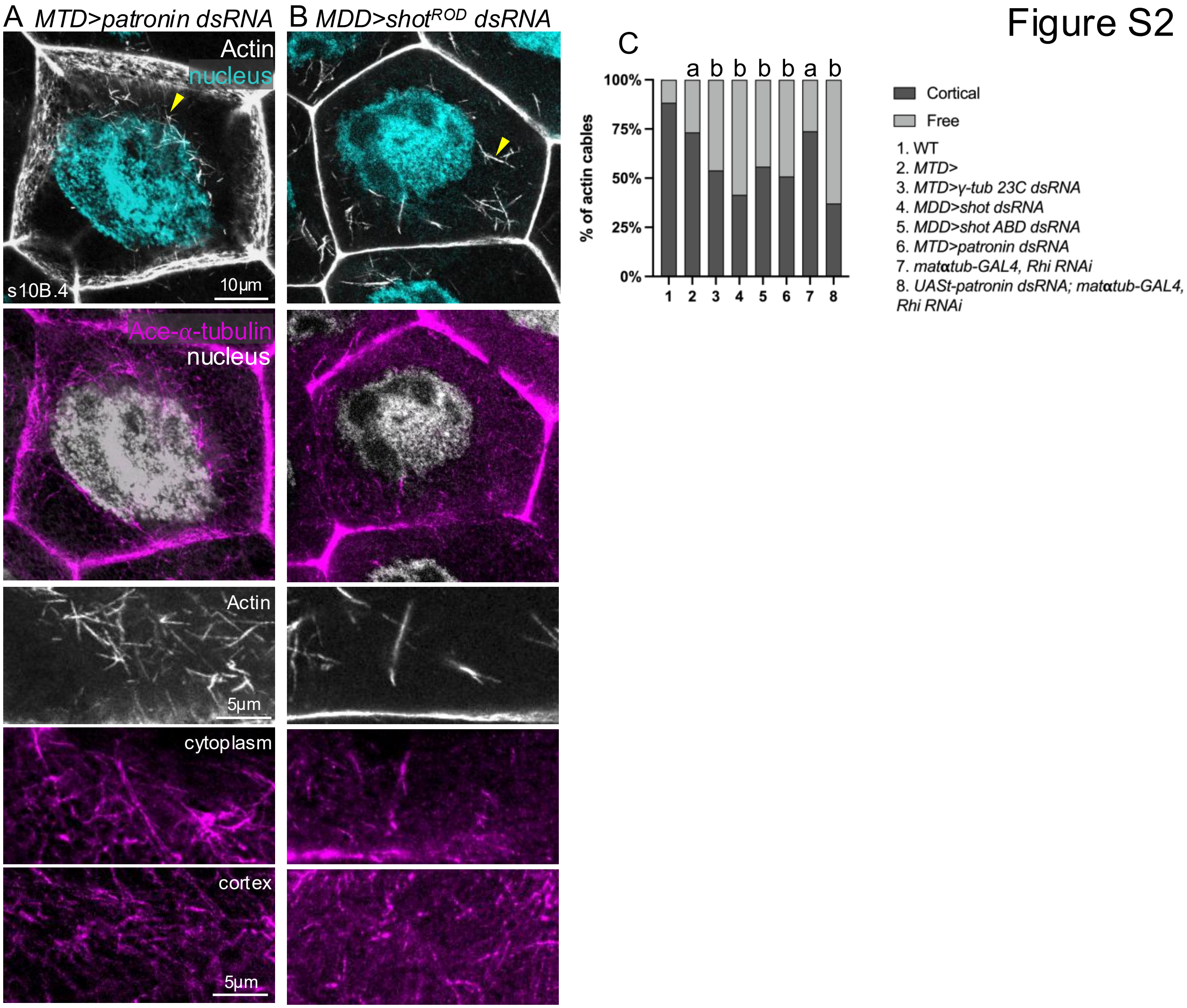
A MTD>patronin dsRNA
B MDD>shot^ROD dsRNA
C
Actin
nucleus
s10B.4
10µm
Ace-α-tubulin
nucleus
Actin
5µm
cytoplasm
cortex
5µm
% of actin cables
100%
75%
50%
25%
0%
1 2 3 4 5 6 7 8
a b b b b a b
Cortical
Free
1. WT
2. MTD>
3. MTD>γ-tub 23C dsRNA
4. MDD>shot dsRNA
5. MDD>shot ABD dsRNA
6. MTD>patronin dsRNA
7. matαtub-GAL4, Rhi RNAi
8. UASt-patronin dsRNA; matαtub-GAL4, Rhi RNAi

### Figure S3

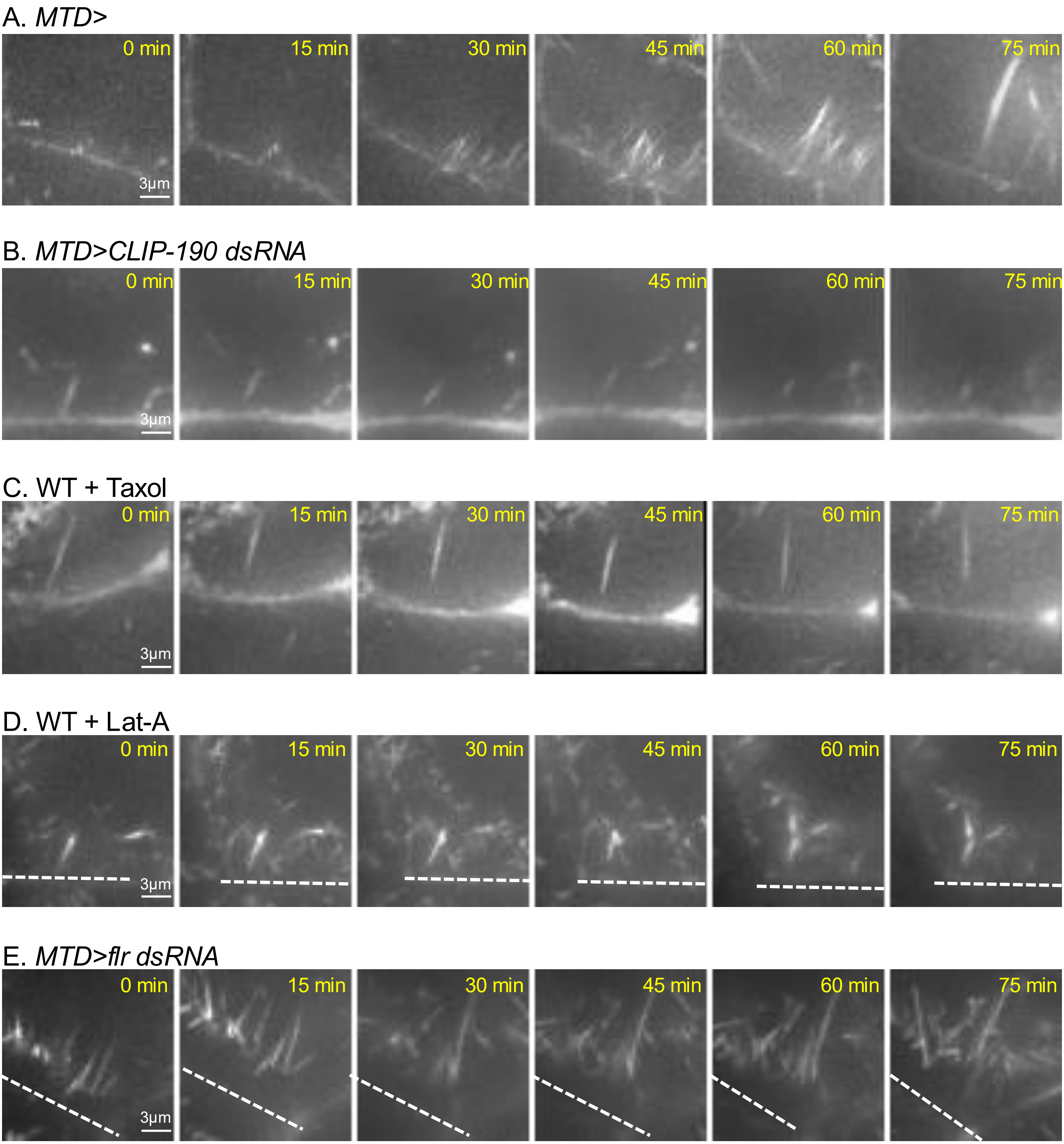

A. *MTD>*
0 min
15 min
30 min
45 min
60 min
75 min
3μm
B. *MTD>CLIP-190 dsRNA*
0 min
15 min
30 min
45 min
60 min
75 min
3μm
C. WT + Taxol
0 min
15 min
30 min
45 min
60 min
75 min
3μm
D. WT + Lat-A
0 min
15 min
30 min
45 min
60 min
75 min
3μm
E. *MTD>flr dsRNA*
0 min
15 min
30 min
45 min
60 min
75 min
3μm

### Figure S4

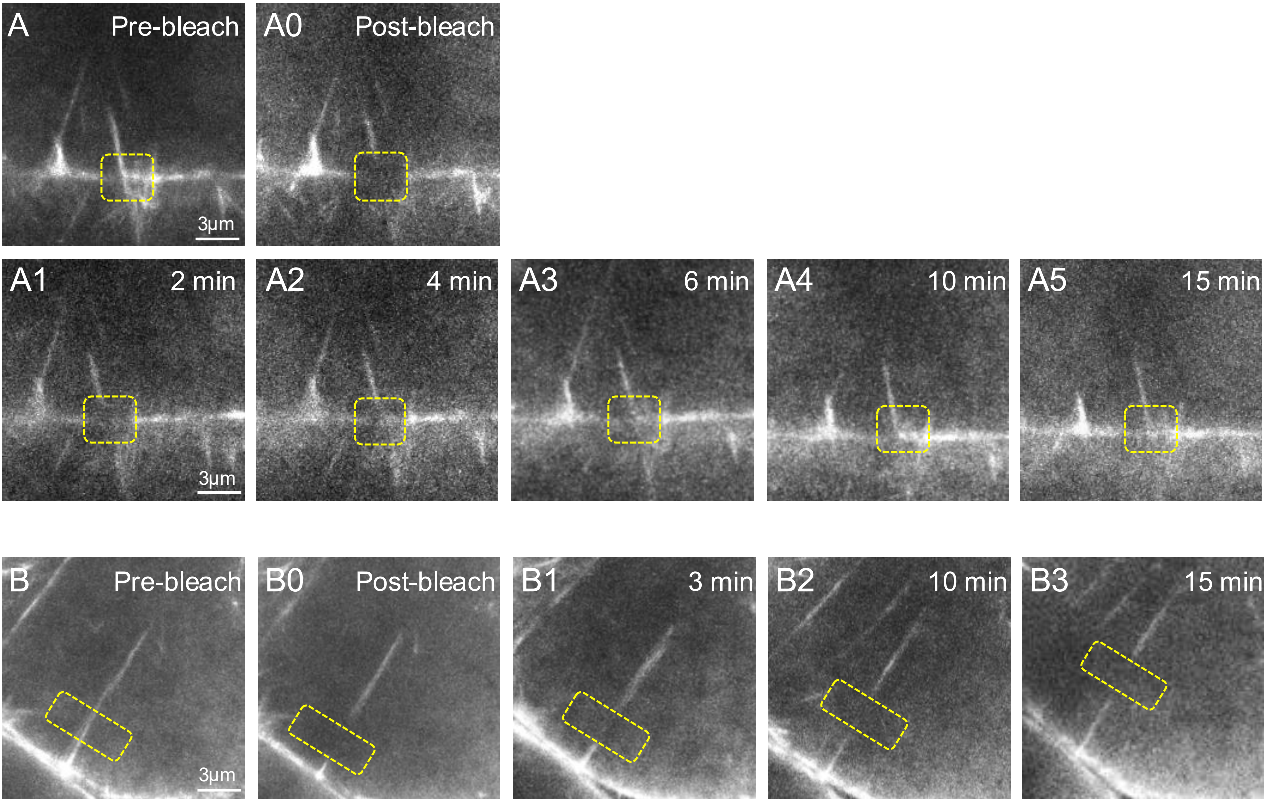

A
Pre-bleach
3μm
A0
Post-bleach
A1
2 min
3μm
A2
4 min
A3
6 min
A4
10 min
A5
15 min
B
Pre-bleach
3μm
B0
Post-bleach
B1
3 min
B2
10 min
B3
15 min

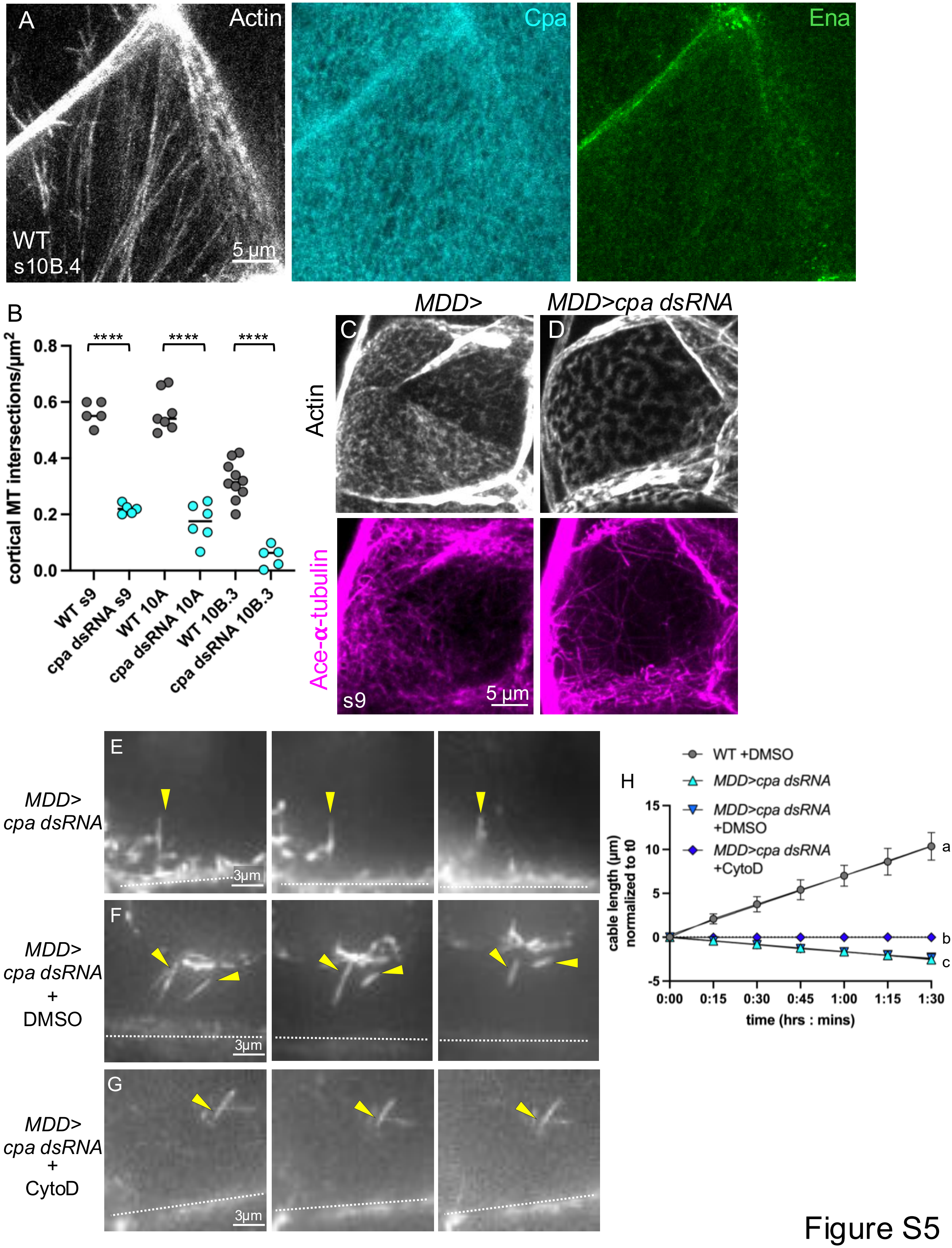

A
Actin
Cpa
Ena
WT
s10B.4
5 µm
B
cortical MT intersections/µm²
0.8
0.6
0.4
0.2
0.0
****
****
****
WT s9
cpa dsRNA s9
WT 10A
cpa dsRNA 10A
WT 10B.3
cpa dsRNA 10B.3
MDD>
MDD>cpa dsRNA
C
D
Actin
Ace-α-tubulin
s9
5 µm
E
MDD> cpa dsRNA
3µm
F
MDD> cpa dsRNA + DMSO
3µm
G
MDD> cpa dsRNA + CytoD
3µm
H
WT +DMSO
MDD>cpa dsRNA
MDD>cpa dsRNA +DMSO
MDD>cpa dsRNA +CytoD
cable length (µm) normalized to t0
15
10
5
0
-5
a
b
c
0:00
0:15
0:30
0:45
1:00
1:15
1:30
time (hrs : mins)


Figure S5
